# The RING Finger E3 Ligase RNF25 Protects DNA Replication Forks Independently of its Canonical Roles in Ubiquitin Signaling

**DOI:** 10.1101/2025.01.09.632184

**Authors:** Lilly F. Chiou, Deepika Jayaprakash, Gaith N. Droby, Xingyuan Zhang, Yang Yang, C. Allie Mills, Thomas S. Webb, Natalie K. Barker, Di Wu, Laura E. Herring, Jessica Bowser, Cyrus Vaziri

## Abstract

The DNA damage response (DDR) mechanisms that allow cells to tolerate DNA replication stress are critically important for genome stability and cell viability. Using an unbiased genetic screen we identify a role for the RING finger E3 ubiquitin ligase RNF25 in promoting DNA replication stress tolerance. In response to DNA replication stress, RNF25-deficient cells generate aberrantly high levels of single-stranded DNA (ssDNA), accumulate in S-phase and show reduced mitotic entry. Using single-molecule DNA fiber analysis, we show that RNF25 protects reversed DNA replication forks generated by the fork remodeler HLTF from nucleolytic degradation by MRE11 and CtIP. Mechanistically, RNF25 interacts with the replication fork protection factor REV7 and recruits REV7 to nascent DNA after replication stress. The role of RNF25 in protecting replication forks is fully separable from its canonical functions in ubiquitin conjugation. This work reveals the RNF25-REV7 signaling axis as an important protective mechanism in cells experiencing replication stress.

## Introduction

DNA replication stress (slowing of DNA synthesis due to various obstacles) poses a threat to all cells. Unresolved DNA replication stress can compromise genome stability or lead to cell death. In the clinic, chemotherapy-induced replication stress can be a valuable strategy for killing cancer cells. The success of chemotherapy depends critically on agents that can induce replication stress and irreparable DNA damage in cancer cells.^1–3^

Common causes of replication stress include bulky DNA lesions, transcription-replication collisions, R-loop formation, nucleotide depletion, or aberrant origin firing.^1–6^ Nucleotide depletion and aberrant origin firing are common features of many cancer cells that likely contribute to the high levels of intrinsic replication stress in tumors.^2,7,8^ DNA replication forks experiencing replication stress are vulnerable and can ‘collapse’ to generate lethal DNA Double Stranded Breaks (DSBs). To mitigate the threat of replication stress, cells have evolved <u>D</u>NA <u>D</u>amage <u>R</u>esponse (DDR) mechanisms that protect and resolve stalled and collapsed DNA replication forks.

For example, specialized trans-lesion synthesis (TLS) DNA polymerases can bypass certain DNA lesions and sustain ongoing DNA replication on damaged templates.^9^ The primase-polymerase PRIMPOL can also restart replication downstream of stalled leading strands, thereby generating a ssDNA gap distal to newly reprimed forks.^10–12^ Fork reversal is another mechanism to manage replication fork stalling. During fork reversal, the leading and lagging nascent DNA strands anneal to form a three-way ‘chicken-foot’ and the undamaged lagging strand may be used as a template for leading strand synthesis.^13,14^ Although fork reversal is a protective mechanism, this process generates a single-ended DSB (seDSB) which is vulnerable to nucleolytic attack.^15–17^

Degradation of reversed forks is prevented by fork protection factors such as BRCA2, FANCD2, ABRO1, 53BP1, and REV7.^18–22^ In the absence of fork protection, reversed DNA replication forks are degraded by a wide variety of nucleases including MRE11, EXO1, DNA2, and CtIP.^15,18–23^ Excessive degradation of reversed forks can generate DSBs and lead to cell death. Therefore, protection of reversed replication forks is a crucial element of replication stress tolerance. Understanding mechanisms of fork protection during replication stress could reveal novel strategies to sensitize cancer cells to intrinsic or therapy-induced DNA replication stress.

Here we sought to identify new mechanisms of replication stress resistance. We focused on E3 ligases as candidate mediators of replication stress resistance for several reasons. E3 ligases play central roles in regulation of DNA replication and repair, both through ubiquitin signaling and through non-catalytic scaffold or chaperone roles.^24–27^ Pathways involving E3 ligases are often rewired pathologically in neoplastic cells and therefore represent appealing targets for cancer therapy.^28,29^ Potent small molecule inhibitors have been developed against several E3 ligases that sustain cancer cells.^30–33^ E3 ligases are under-utilized as therapeutic targets and there remain many attractive opportunities for E3 ligase-directed small molecules and therapies. Moreover, there are many orphan E3 ligases with no known function. Therefore, opportunities exist to discover new E3 ligase-driven mechanisms that sustain cancer cells. Here, we identify a new role for the understudied E3 ubiquitin ligase, RNF25, in conferring replication stress tolerance. We demonstrate that RNF25 interacts with and recruits another fork protection factor REV7 to replication forks. Notably, ubiquitin conjugating activity is dispensable for RNF25-mediated fork protection. In the absence of RNF25, replication forks are extensively degraded by the nucleases MRE11 and CtIP. Taken together, our work establishes an important mechanism of DNA replication fork protection involving RNF25.

## Results

### CRISPR screening reveals RING finger E3 ubiquitin/SUMO ligase genes required for WEE1i-tolerance

We constructed a sgRNA library targeting 317 RING finger-E3 ubiquitin and SUMO ligases (∼10 unique sgRNAs per gene) also including 1000 non-targeting (NT) control sgRNAs. Using our sgRNA library we performed a CRISPR-Cas9 ‘dropout’ screen to identify E3 ligase dependencies of Pa02C and Pa03C Pancreatic Ductal Adenocarcinoma (PDAC) cell lines cultured in the absence and presence of the WEE1 inhibitor (WEE1i) AZD-1775.

We used WEE1 inhibition as a source of replication stress for several reasons: Mechanistically, WEE1 inhibition induces DNA replication stress via CDK2 over-activation,^34^ which is a feature of many cancer cells.^35^ Thus, results identified with WEE1i treatments are broadly relevant to cancer. WEE1 inhibitors are emerging as attractive drugs for treatment of several malignancies.^36–43^ Furthermore, there is overlap between mechanisms of action of WEE1i and many other anti-cancer drugs used as single agents and in combination therapies.^4^ Therefore, targeting mediators of WEE1i-resistance might also improve the efficacy of other agents that also cause replication stress. We performed our screens using Pa02C and Pa03C cell lines because PDAC is being evaluated for responsiveness to WEE1i (Clinical Trials Identifier NCT06015659). Pa02C and Pa03C cell lines are well characterized and have genetic features that are highly representative of PDAC patient tumors,^44,45^ including KRAS mutants that are known to induce DNA replication stress.^46^

The workflow of our CRISPR-Cas9 screening platform is summarized in Fig. 1a. We sequenced the genomes of cells that were cultured +/- WEE1i, and used the Völundr computational pipeline^47^ to quantify all sgRNAs remaining after 20 population doublings (PDs). The volcano plots in Fig. 1b, c depict the relative loss or enrichment of different sgRNAs and the statistical significance of those changes (Kolmogorov–Smirnov test) across our test conditions. Gene-targeting sgRNAs were depleted more than NT control sgRNAs in Pa02C and Pa03C cells cultured both with and without WEE1 inhibitor, indicating a broad dependency on E3 ligases for basal survival (Supplementary Fig. 1a, b). For example, sgRNAs targeting essential genes such as *UBE2I*, *TRAIP* and *RNF8* showed significant drop-out both basally and in WEE1i-treated cultures (Supplementary Table 1-4). However, some sgRNAs dropped out more significantly under WEE1i treatment conditions (see highlighted sgRNAs in Fig. 1b, c). Taken together, our CRISPR-Cas9 screens identified E3 ligases that are required for basal growth of PDAC cells and for tolerating WEE1i-induced replication stress.

**Fig. 1:**
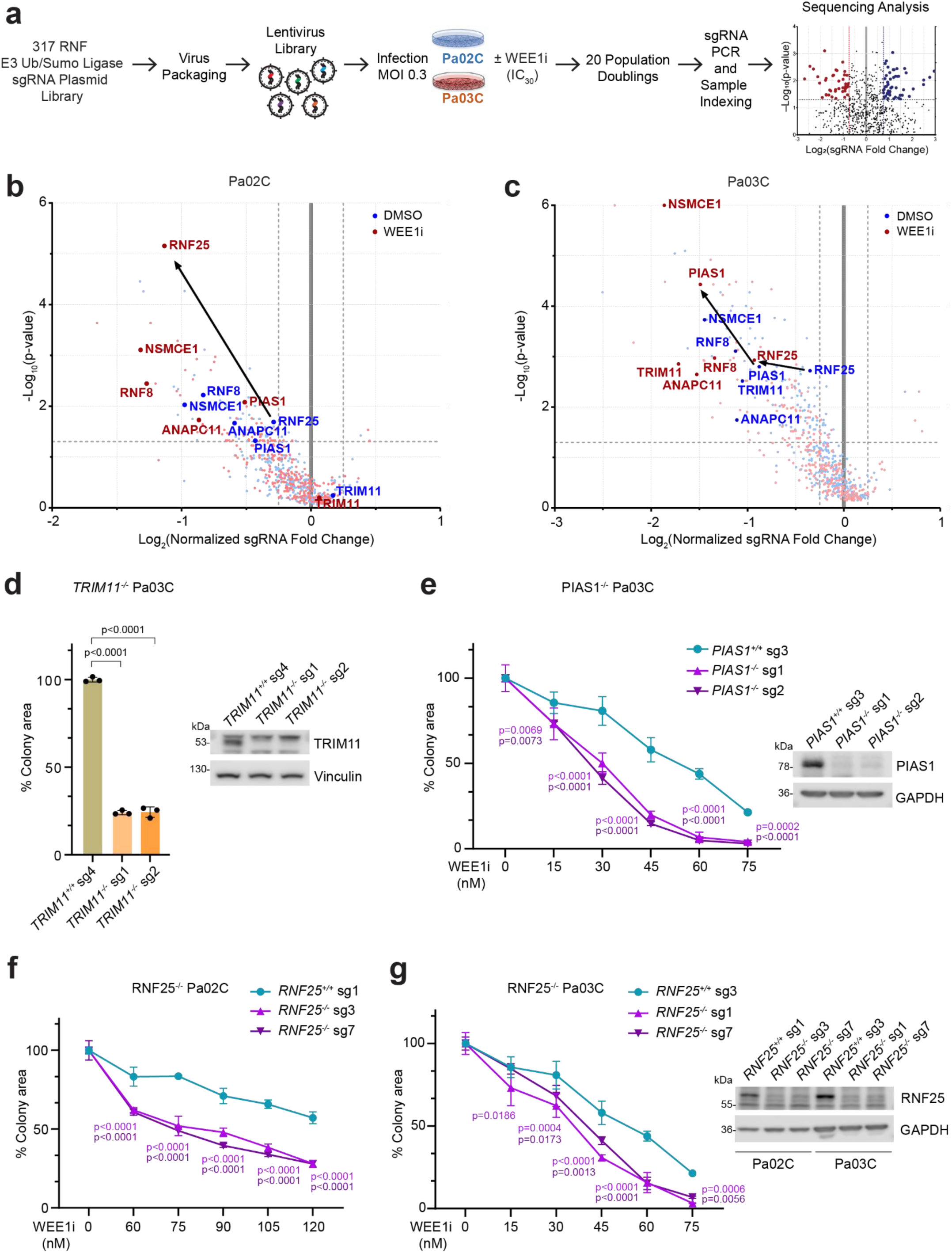
CRISPR screening reveals RING finger E3 ubiquitin/SUMO ligase genes required for WEE1i-tolerance. **a** Experimental design of the CRISPR screens performed in this study. **b, c** Results of CRISPR screens showing dropout of sgRNAs in vehicle (DMSO)- and WEE1i-treated Pa02C and Pa03C cells. Log_2_-transformed sgRNA abundance scores and statistical significance were calculated using Völundr. **d** Effect of TRIM11-directed sgRNAs on clonogenic survival of Pa03C cells. **e** Effect of PIAS-deletion on WEE1-sensitivity in Pa03C cells. **f, g** Effect of RNF25-deficiency on WEE1-sensitivity in Pa02C (f) and Pa03C (g) cells. All data in (d-g) represent mean ± SD from triplicate wells and are representative of two biological replicates. Statistics: two-way ANOVA with Tukey’s multiple comparisons test.

### RNF25 and PIAS1 are necessary for WEE1i-tolerance

We independently validated some of the E3 ligase dependencies identified by our CRISPR dropout screens. We selected *TRIM11*, *RNF25* and *PIAS1* for validation because based on the DEPMAP portal these genes are non-essential for most cancer cell lines. As shown in Fig. 1c and Supplementary Table 3, 4, sgRNAs targeting *TRIM11* were depleted from Pa03C cells (even in the absence of WEE1i treatment), indicating that *TRIM11* is important for basal survival. Consistent with this result, acute ablation of *TRIM11* using two independent sgRNAs led to ∼80% decrease in clonogenic survival of Pa03C cells when compared with Pa03C cells expressing non-targeting (NT) sgRNA (Fig. 1d). We conclude that *TRIM11* is necessary for proliferation of Pa03C cells. Our findings are also consistent with a recent report showing that *TRIM11* is required for survival of PDAC both in vitro and in vivo.^48^

Figure 1c and Supplementary Table 3, 4 also show that dropout of *PIAS1*-directed sgRNAs in Pa03C cells was increased with WEE1i treatment. Consistent with this result, two independent polyclonal *PIAS1^-/-^* Pa03C cell lines (generated using different sgRNAs) were WEE1i-sensitive when compared with control *PIAS1^+/+^*Pa03C cells expressing non-targeting (NT) sgRNA (Fig. 1e). Therefore, *PIAS1* is necessary for WEE1i-tolerance of Pa03C cells. In our CRISPR screens, sgRNAs targeting *RNF25* showed increased dropout after WEE1i treatment in both Pa02C and Pa03C cells (Fig. 1b, c and Supplementary Table 1-4). In validation experiments, *RNF25^-/-^*Pa02C and Pa03C cells (generated using two independent sgRNAs) were WEE1i-sensitive when compared with *RNF25^+/+^* control cells (Fig. 1f, g). Additionally, based on rank orders of dropout, sgRNAs targeting RNF25 and PIAS1 were depleted more heavily in transformed and aggressive PDAC cell lines when compared with hTERT-immortalized but non-transformed normal pancreatic epithelial cells (designated hTERT-HPNE – see Supplementary Fig. 1c-g). Taken together, the results of Fig. 1d-g validate our CRISPR screen and identify specific new roles for PIAS1 and RNF25 in tolerating WEE1 inhibition.

### RNF25 prevents ssDNA accumulation and facilitates S-phase and G2/M progression of WEE1i-treated cells

We hypothesized that RNF25 and PIAS1 might suppress accumulation of DNA damage in cells experiencing WEE1i-induced DNA replication stress. Consistent with this hypothesis, *RNF25^-/-^* Pa03C cells aberrantly accumulated excessive levels of the ssDNA marker phospho-RPA (pRPA) when compared with *RNF25^+/+^* controls (Fig. 2a). Aberrantly high levels of pRPA (up to 2.50-fold higher peak expression) were detectable at 24 h, 48 h, 72 h, and 13 days after WEE1i-treatment in *RNF25^-/-^* cells (Fig. 2a). After 13 days of WEE1i treatment, we also observed high (2.73-fold) and persistent levels of the DSB marker phospho-ATM (S1981) in *RNF25^-/-^* cells when compared with *RNF25^+/+^* control cultures (Fig. 2a). Taken together, our results indicate that RNF25 averts formation of excessive ssDNA and DSB in WEE1i-treated cells. WEE1i-treated *RNF25^-/-^* cells accumulated aberrantly in S-phase and G2/M when compared with *RNF25^+/+^* cells (Fig. 2b). Using flow-cytometry-based measurements we found that *RNF25^-/-^* cells in mitosis contained high levels of ssDNA after WEE1i treatment when compared with parental *RNF25^+/+^* cells (Fig. 2c). We conclude that in the absence of RNF25, WEE1i-treated cells accumulate aberrantly high levels of ssDNA during DNA replication that leads to delays in S-phase and G2/M progression.

**Fig. 2:**
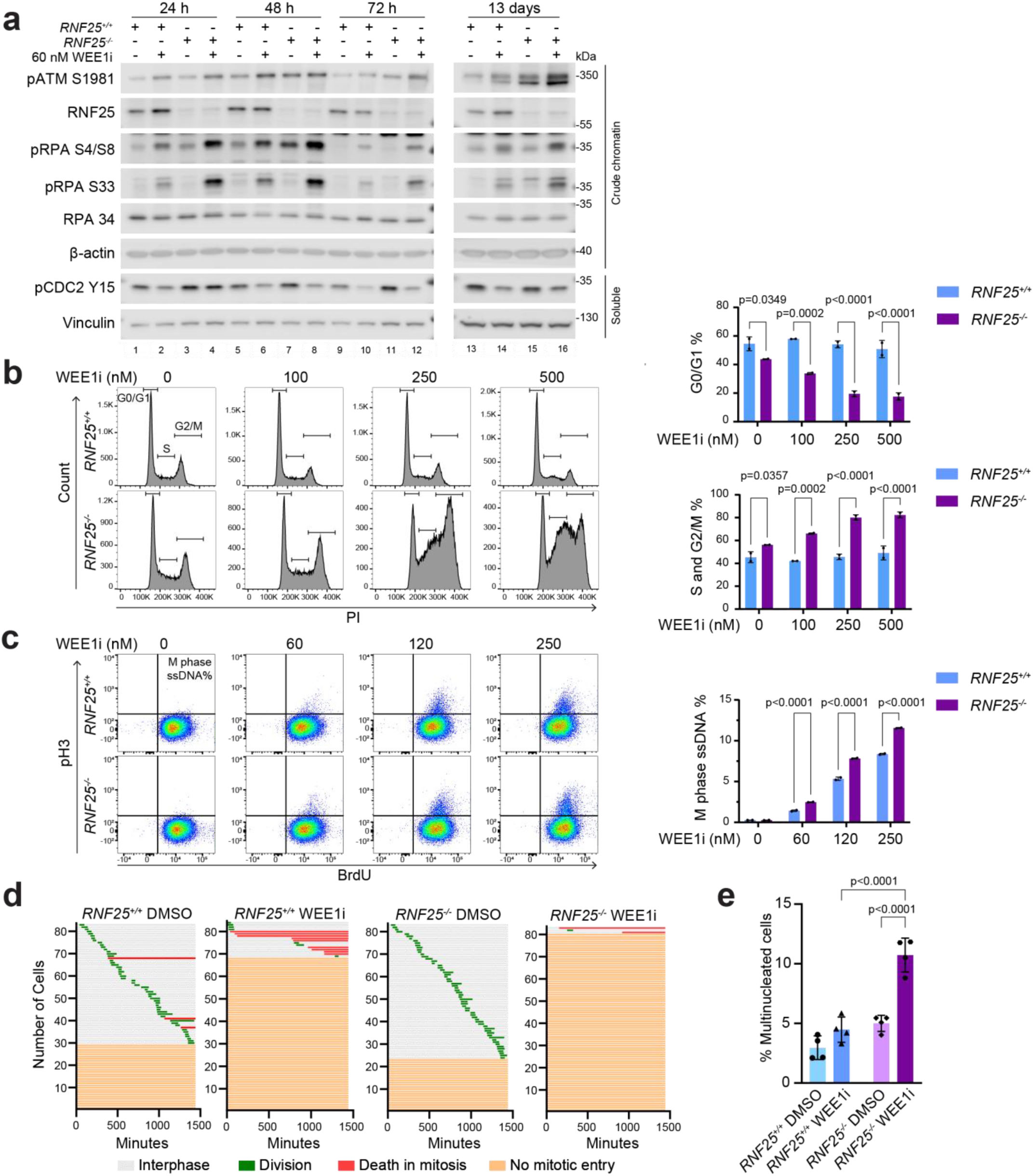
RNF25 prevents ssDNA accumulation and facilitates S-phase and G2/M progression of WEE1i-treated cells. **a** Effect of WEE1i treatment on levels of the indicated DNA damage markers in *RNF25^+/+^* and *RNF25^-/-^* Pa03C cells. **b** Effect of WEE1i treatment on cell cycle profiles of *RNF25^+/+^* and *RNF25^-/-^* Pa03C cells. **c** Effect of WEE1i treatment on levels of ssDNA in pH3-postive (mitotic) nuclei in *RNF25^+/+^* and *RNF25^-/-^* Pa03C cells. **d** Cell fate maps showing effect of WEE1i-treatment on interphase and mitotic progression of *RNF25^+/+^* and *RNF25^-/-^* Pa03C cells. **e** Effect of WEE1i-treatment on multinucleation in *RNF25^+/+^* and *RNF25^-/-^* Pa03C cells. All data represent mean ± SD from four focus locations. Ordinary one-way ANOVA with Tukey’s multiple comparisons test was performed to analyze statistical significance. All flow cytometry data represent mean ± SD from duplicate dishes and are representative of two biological replicates. Statistics: two-way ANOVA with Sidak’s multiple comparisons test.

To formally test the hypothesis that RNF25 is required for normal mitotic progression of WEE1i-treated cells, we performed live cell imaging and generated mitotic fate maps of *RNF25^+/+^* and *RNF25^-/-^*cultures growing in the absence or presence of WEE1i. As shown in Fig. 2d, the cell cycle intervals and mitotic fates of *RNF25^+/+^* and *RNF25^-/-^* cells were not significantly different in the absence of WEE1i. However, *RNF25^-/-^* cells were more sensitive to WEE1i-induced inhibition of mitotic entry when compared with *RNF25^+/+^*cells (Fig. 2d). Moreover, after WEE1i treatment, the percentage of multinucleated cells was significantly higher in *RNF25^-/-^* cells when compared to the *RNF25^+/+^*controls (Fig. 2e). Therefore, we conclude that RNF25 sustains progression of WEE1i-treated cells through S-phase and into G2/M and prevents mitotic defects.

### RNF25-dependent WEE1i-tolerance is unrelated to NF-κB, ERK, or translation quality control

RNF25 has previously been implicated in suppression of NF-κB signaling and reactivation of ERK during chronic drug treatment.^49,50^ However, under our standard experimental conditions NF-κB activity and ERK1/2 phosphorylation were unaffected by RNF25 status (Supplementary Fig. 2a-d). Therefore, we do not attribute the WEE1i-sensitivity of *RNF25^-/-^* cells to changes in NF-κB or ERK signaling.

RNF25 was also recently shown to cooperate with the E3 ubiquitin ligase RNF14 to mediate ubiquitination and degradation of eEF1A and eRF1 on stalled ribosomes.^51,52^ We considered the possibility that the role of RNF25 in WEE1i-tolerance might be related to its function in a ribosome collision sensing pathway. However, RNF14 was dispensable for WEE1i tolerance (Supplementary Fig. 2e), indicating that WEE1 inhibition does not create dependency on ribosome quality control. In DNA repair-compromised PDAC cells, persistent nuclear DNA damage generates cytosolic DNA that activates the cGAS-STING pathway.^53^ As shown in Supplementary Fig. 2f, we observed increased levels of phospho-IRF3 (S386) and phospho-STING (S366), markers of cGAS-STING signaling in *RNF25^-/-^*cells after WEE1i treatment. Taken together, our results suggest that the role of RNF25 in promoting WEE1i-tolerance is related to genome maintenance, and separable from its functions in ERK signaling, NFκB activation and translation quality control.

### PIAS1 promotes S-phase and G2/M progression in WEE1i-treated PDAC cells

We also defined the effect of PIAS1-deficiency on cell cycle responses to WEE1i. As expected, *PIAS1^-/-^* cells (lacking Protein Inhibitor of Activated STAT1) contained higher levels of phosphorylated pSTAT1 (Tyr701) when compared with control *PIAS1^+/+^* cells (Supplementary Fig. 3a). However, STAT1-depletion (using siRNA) did not affect clonogenic survival of WEE1i-treated cells (Supplementary Fig. 3b). Therefore, the WEE1i-sensitivity of *PIAS1^-/-^* cells cannot be attributed to excessive STAT1 signaling. Similar to *RNF25*-deficient cells, *PIAS1^-/-^* Pa03C cells treated with WEE1i aberrantly accumulated high levels of the ssDNA marker phospho-RPA (pRPA) when compared with *PIAS1^+/+^* control cultures (Supplementary Fig. 3a). We also detected higher levels of S1981-phosphorylated-ATM in WEE1i-treated *PIAS1^-/-^* cells when compared with *PIAS1^+/+^*controls (Supplementary Fig. 3a), indicating that PIAS1 averts DSB formation. After WEE1i-treatment, *PIAS1^-/-^* cells accumulated aberrantly in S- and G2/M-phase when compared with *PIAS1^+/+^*cells (Supplementary Fig. 3c). In live cell imaging experiments, cell cycle intervals and mitotic fates of *PIAS1^+/+^*and *PIAS1^-/-^* cells were similar in the absence of WEE1i (Supplementary Fig. 3d). However, after WEE1 inhibition, *PIAS1^-/-^* cells were significantly more prone to death in mitosis when compared with *PIAS1^+/+^* cells (Supplementary Fig. 3d, e). We conclude that similar to RNF25, PIAS1 prevents acquisition of persistent replication-associated DNA damage and promotes survival in response to WEE1i-induced genotoxicity.

Roles of PIAS1 in DNA replication and DDR signaling are established^54,55^ yet there is no known mechanism that explains how RNF25 promotes tolerance of DNA replication stress. Therefore, we used DNA fiber assays to determine with high resolution how RNF25-deficiency impacts DNA replication dynamics. Using DNA fiber assays, *RNF25*^-/-^ Pa02C and Pa03C cells both showed reduced rates of DNA replication fork movement when compared with their respective *RNF25^+/+^*parental cells, even without pharmacologically-induced DNA replication stress (Fig. 3a). Similar results were obtained in H1299 lung adenocarcinoma cells when using two independent *RNF25^-/-^* knockout clones and two independent siRNAs to transiently deplete RNF25 (Fig. 3c), suggesting that RNF25 promotes tolerance of intrinsically-arising DNA replication stress by promoting DNA replication fork movement. Replication fork reversal is a protective mechanism in response to replication stress, however, the regressed arm of DNA is susceptible to digestion by nucleases leading to fork degradation.^14^ Using the DNA fiber assay, *RNF25^-/-^* Pa02C and Pa03C cells both showed increased levels of fork degradation (Fig. 3b). The same phenotype was observed in H1299 cells lacking RNF25 (Fig. 3d). Therefore, we conclude that RNF25 promotes both DNA replication fork movement as well as replication fork protection in response to replication stress.

**Fig. 3:**
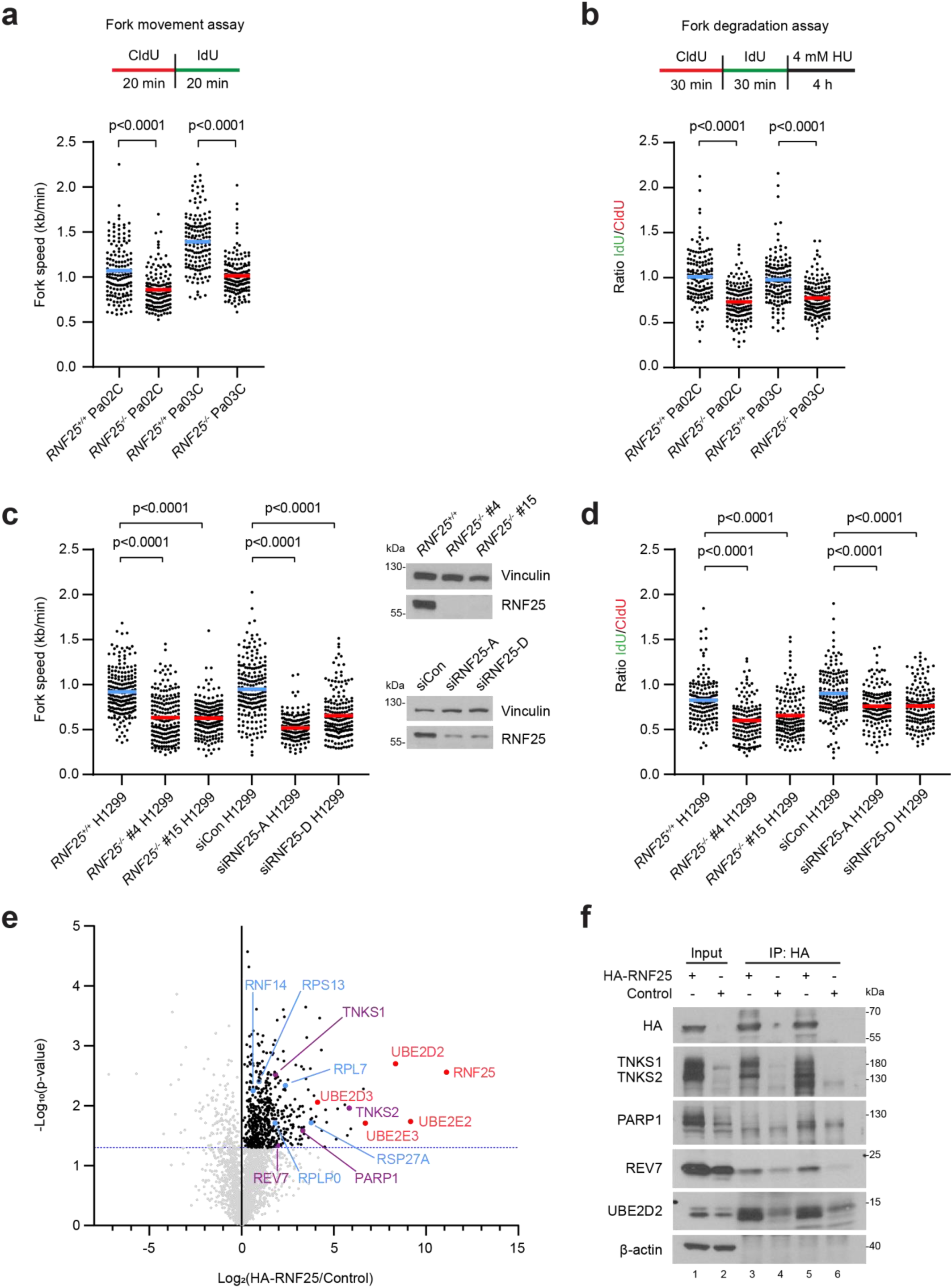
RNF25 mediates fork movement and fork protection. **a** Top: Schematic of fork movement assays. Bottom: *RNF25^+/+^* and *RNF25^-/-^* Pa02C and Pa03C cells were labeled with CldU and IdU and fork speed was calculated by dividing tract lengths by total pulse time. **b** Top: Schematic of fork degradation assays. Bottom: *RNF25^+/+^* and RNF25^-/-^ Pa02C and Pa03C cells were labeled with CldU and IdU, then treated with HU to induce fork stalling. The ratio of IdU to CldU tracts was used as a readout of fork degradation. **c, d** The fork movement assay (c) and fork degradation assay (d) were performed in *RNF25^+/+^* H1299 cells and two independent *RNF25^-/-^* clones, as well as *RNF25^+/+^* H1299 cells treated with siControl, and two independent siRNAs targeting RNF25. **e** Volcano plot of the HA-RNF25 IP-MS results in H1299 cells. Log_2_ fold change ratios (HA-RNF25/Control) on the x-axis and -log_10_ p-values on the y-axis. -log_10_ p-value > 1.3 indicates statistical significance (two-tailed Student’s t-test). Red: RNF25 and E2 ligases; Blue: known RNF25-interacting factors involved in translation quality control; Purple: proteins involved in DNA replication. **f** H1299 cells expressing HA-RNF25 were immunoprecipitated using anti-HA beads. Lanes 3 and 4 were immunoprecipitated using anti-HA beads from Thermo Fisher Scientific. Lanes 5 and 6 were immunoprecipitated using anti-HA beads from MBL. All DNA fiber assays shown (a-d) are a representative experiment of two biological replicates. 150 fibers were quantified per condition in a, b, d; 200 fibers were quantified per condition in c. Means are indicated with bars. Statistics: two-tailed Mann-Whitney test.

### RNF25 associates with DNA replication fork protection factors

To identify mediators and mechanisms of RNF25-dependent DNA replication fork progression, we immunopurified HA-RNF25 from cultured cells and identified its binding partners using proteomics-based mass spectrometry. These experiments were necessarily performed in H1299 cells in which we were able to achieve higher HA-RNF25 expression levels when compared with PDAC cell lines. We identified the known E2 enzymes for RNF25, UBE2D2 and UBE2E3,^56^ as well as other E2 enzymes within this family, UBE2E2 and UBE2D3 (Fig. 3e). Known RNF25-interacting factors involved in translation quality control (including RPS27A, RPLP0, RPL7, RPS13, and RNF14)^51^ were also present in HA-RNF25 complexes (Fig. 3e). Interestingly, we also identified proteins with known roles in DNA replication in the RNF25 complex including PARP1, REV7, TNKS1 (PARP5A), and TNKS2 (PARP5B).^22,57–63^ Using independent co-immunoprecipitation experiments we validated the interactions of RNF25 with PARP1, TNKS1/2 and REV7 (Fig. 3f). Levels of TNKS1/2 and REV7 were increased in response to ectopically-expressed HA-RNF25 (Fig. 3f), potentially suggesting that RNF25 stabilizes these binding partners.

### RNF25 localizes to sites of DNA replication fork stalling and promotes recovery from replication stress

In HU-treated H1299 cells we observed a 2.36-fold increase in the levels of chromatin-bound RNF25 (peaking at 1 h post-recovery), suggesting that RNF25 is recruited to sites of replication stalling (Fig. 4a, b, top panels). In response to DNA replication stress, multiple overlapping and redundant DNA repair processes cooperate to promote fork recovery, including TLS, repriming and fork reversal. Therefore, we asked whether there was compensatory activation of those DNA repair pathways in *RNF25*-deficient cells both during, and after recovery from HU-induced replicative arrest. As shown in Fig. 4a, in *RNF25^-/-^* cells treated with HU we observed up to 2.88-fold increase in levels of PCNA-mono-ubiquitylation (and concomitant increases in levels of chromatin-bound RAD18, REV1, and Pol η (3.03-fold, 3.27-fold, and 4.49-fold increases respectively)) when compared with HU-treated *RNF25^+/+^* cells. Therefore the TLS pathway is hyperactivated in the absence of RNF25. Levels of chromatin-bound HLTF and ZRANB3 were also more abundant (2.30-fold and 1.51-fold respectively) in HU-treated *RNF25^-/-^* cells when compared with *RNF25^+/+^* cells, also indicating increased activation of fork reversal in the absence of RNF25. The levels of chromatin-bound FANCD2 (a reversed fork protection factor) were increased by up to 2.01-fold in HU-treated *RNF25^-/-^* cells when compared with wild-type cells (Fig. 4a), also consistent with increased fork reversal when RNF25 is absent. We also observed up to 2.09-fold increase in levels of chromatin-bound primase-polymerase PRIMPOL in HU-treated *RNF25^-/-^*cells when compared with parental *RNF25^+/+^* cells, suggesting that repriming is also stimulated to promote fork recovery when RNF25 is absent. Qualitatively similar results were obtained in two *RNF25^-/-^*clones (Fig. 4a) and when we depleted RNF25 using two independent siRNAs (Fig. 4b). Taken together, these results suggest that there is compensatory activation of TLS, repriming and fork reversal in HU-treated cells lacking RNF25. However, in response to HU treatment, RNF25-deficient cells accumulated high levels of γH2AX-ubiquitylation (3.27-fold increase) when compared to RNF25-replete cells (Fig. 4a, b). Therefore, despite compensatory increases in activation of other stress tolerance pathways, RNF25-deficient cells likely accumulate persistent DSB and fail to recover normally from HU-induced DNA replication stress.

**Fig. 4:**
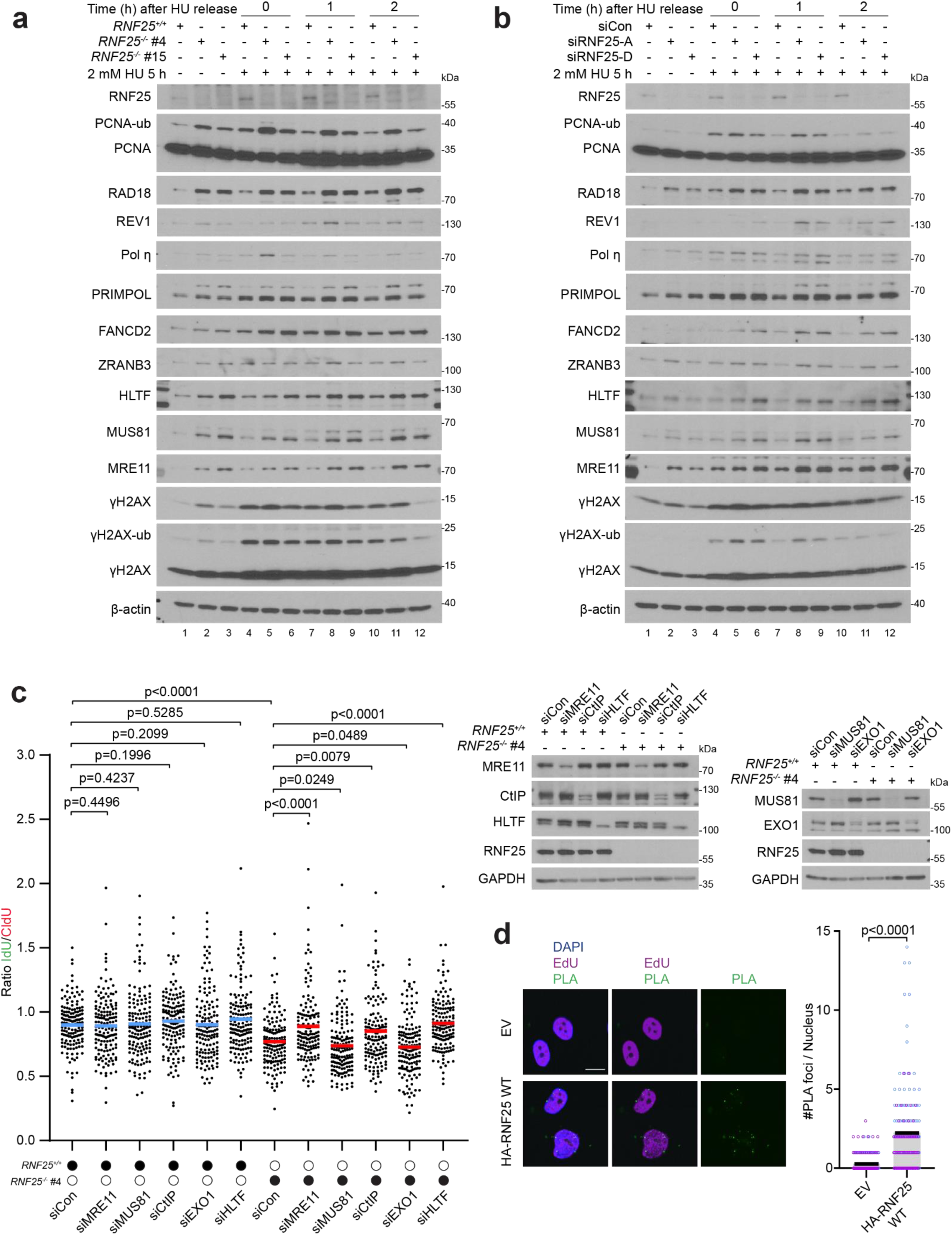
RNF25 localizes to sites of DNA replication fork stalling and promotes recovery from replication stress. **a** *RNF25^+/+^* and two independent *RNF25^-/-^* H1299 clones were treated with 2 mM HU for 5 h, then washed and timepoints taken at 0, 1, and 2 h after release from HU for analysis by immunoblotting. **b** *RNF25^+/+^* H1299 cells were transfected with control siRNA and two independent siRNAs targeting RNF25 for 48 h, before being assayed as described in (a). **c** *RNF25^+/+^* and *RNF25^-/-^* H1299 cells were transfected with indicated siRNAs for 48 h before conducting the DNA fiber fork degradation assay. A total of 150 fibers were measured per condition and statistics analyzed using a two-tailed Mann-Whitney test. Means are indicated with bars. This figure is a representative experiment from two biological replicates. **d** SIRF assay between HA-RNF25 and EdU in *RNF25^-/-^* H1299 cells. Left: representative images of foci formation; scale bar is 10 μm. Right: the number of foci per nucleus was quantified using ImageJ. Each biological replicate is shown in blue and pink, with means indicated with bars. Number of nuclei quantified per condition (replicate 1-blue, replicate 2-pink): EV: (84, 79); HA-RNF25 WT: (90, 75). Statistics: two-tailed Mann-Whitney test.

Failure to recover from DNA replication stress can be caused by aberrant nucleolytic degradation of reversed replication forks.^15^ We noticed that following HU treatment, the chromatin-binding of nucleases MRE11 and MUS81 was increased (by up to 4.66-fold and 3.34-fold) in RNF25-deficient cells when compared with RNF25-expressing controls (Fig. 4a, b). These results suggested that RNF25 may play a role in protecting reversed forks.

To test the hypothesis that RNF25 prevents excessive fork degradation, we used siRNA to deplete a panel of nucleases in *RNF25^+/+^* and *RNF25^-/-^* H1299 cells, then used DNA fiber assays to measure the stability of reversed forks. As shown in Fig. 4c, depletion of MRE11 and CtIP suppressed fork degradation in *RNF25^-/-^* cells, restoring fork stability to levels that were comparable to those in *RNF25^+/+^* cells (Fig. 4c). Knockdown of nucleases MUS81 and EXO1 did not rescue the fork-degradation phenotype of *RNF25^-/-^* cells (Fig. 4c). This suggests that RNF25 protects reversed forks from degradation by a specific subset of nucleases. Importantly, depleting the fork reversal factor HLTF completely rescued the fork degradation phenotype of RNF25-deficient cells, suggesting that the role of RNF25 in fork protection is dependent on the formation of a reversed fork structure (Fig. 4c).

Because we observed increased levels of chromatin-bound RNF25 following HU treatment (Fig. 4a, b), we asked whether RNF25 resides at active DNA replication forks. We used the *in situ* analysis of protein interactions at DNA replication forks (SIRF) assay^64^ to measure proximity between RNF25 and nascent DNA. We used a doxycycline-inducible vector to complement *RNF25^-/-^* H1299 cells with HA-RNF25 (Supplementary Fig. 4a). Ectopically-expressed HA-RNF25 was expressed at a level similar to that of endogenous RNF25 in the parental *RNF25^+/+^*H1299 cells. We observed significantly increased numbers of PLA foci in HA-RNF25 expressing cells when compared with cells transduced with empty vector (Fig. 4d). These results suggest that RNF25 is present at nascent DNA during basal replication.

### Structural basis of RNF25-mediated replication fork protection

To define RNF25 domains that are important for mediating fork protection we complemented *RNF25^-/-^* cells with a panel of HA-RNF25 variants harboring inactivating mutations in key functional domains (identified by previous studies^56,65^). Figure 5a summarizes the HA-RNF25 mutants we used for our reconstitution experiments. We also generated point mutations C135/138S in the RING domain of RNF25 that abolish binding to E2 ubiquitin-conjugating enzymes.^65^ Additionally, the RNF25 RING domain contains a conserved motif which has been shown to mediate binding with poly-ADP ribose (PAR) chains.^66^ Because our IP-MS experiments revealed PARPs as RNF25-binding partners, we also generated C198/201A point mutations in RNF25 to disrupt its interactions with auto-PARylated PARPs.

**Fig. 5:**
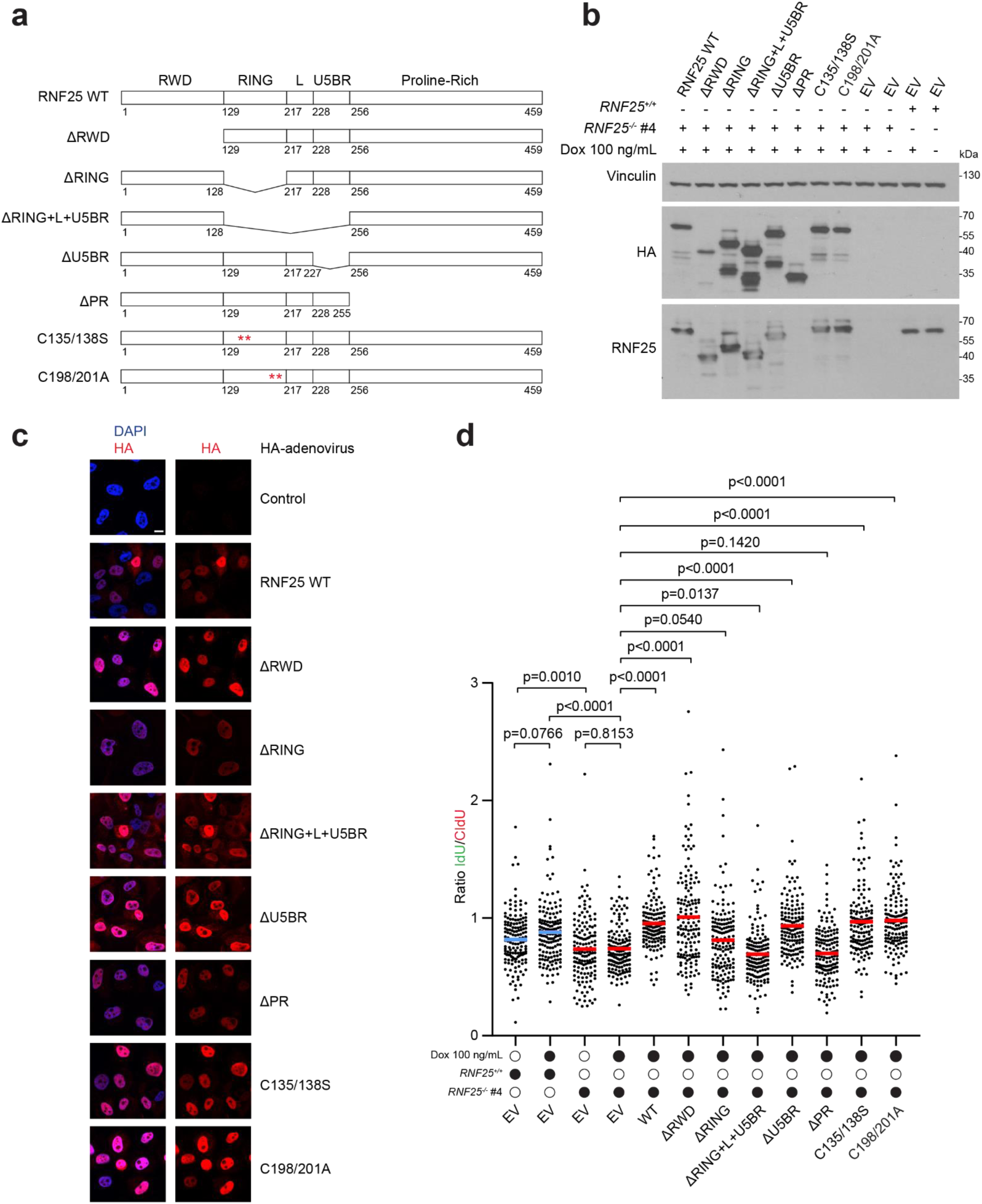
Structural basis of RNF25-mediated replication fork protection. **a** Schematic of full-length human RNF25 WT and RNF25 mutants used in this study. **b** RNF25 WT and mutants from (a) were constructed with an N-terminal HA-tag in a doxycycline-inducible vector and expressed in *RNF25^-/-^* H1299 cells after incubation with doxycycline for 48 h. The RNF25 antibody used recognizes a fragment of RNF25 between residues 409-459, therefore the ΔPR fragment is not recognized in this blot. **c** Immunofluorescence of adenovirally expressed N-terminal HA-tagged RNF25 WT and domain mutants in *RNF25^-/-^* H1299 cells. Scale bar is 10 μm. **d** Fork degradation DNA fiber assay in *RNF25^-/-^* H1299 cells expressing RNF25 WT and domain mutants. 150 fibers were analyzed per condition, with means indicated with bars. This figure is a representative experiment from two biological replicates. Statistics: two-tailed Mann-Whitney test.

Expression levels of wildtype and mutant HA-RNF25 variants in the reconstituted *RNF25^-/-^* cells were similar to that of endogenous RNF25 in parental H1299 cells (Fig. 5b). From immunofluorescence microscopy experiments, all HA-RNF25 variants in our panel correctly localized to the nucleus (Fig. 5c). We performed DNA fiber assays to measure the stability of HU-reversed forks in the panel of HA-RNF25-complemented cells. As expected, *RNF25^-/-^*cells transduced with empty vector showed increased fork degradation when compared with parental *RNF25^+/+^* H1299 cells (Fig. 5d). The fork degradation defect of *RNF25^-/-^* cells was fully rescued by doxycycline-inducible expression of wild-type HA-RNF25. Doxycycline had no significant effect on fork degradation in empty vector cells (Fig. 5d). HA-RNF25 mutants lacking the RWD and U5BR domains also fully rescued the fork degradation defects of *RNF25^-/-^* cells, indicating that these domains are dispensable for mediating fork protection. However, the HA-RNF25 mutant lacking the RING domain only partially rescued the fork degradation phenotype of *RNF25^-/-^* cells. The HA-RNF25 mutant lacking the RING finger, the Linker-domain, and U5BR domain, and the HA-RNF25 mutant lacking the Proline-Rich domain, both completely failed to rescue the fork degradation phenotype (Fig. 5d). Interestingly, the HA-RNF25 mutants harboring C198/201A or C135/138S substitutions both fully rescued the fork degradation phenotype of *RNF25^-/-^*cells. Therefore, associations of RNF25 with PAR and E2 ubiquitin-conjugating enzymes are dispensable for fork protection. We conclude that RNF25 plays a non-catalytic role in protecting DNA replication forks.

### RNF25 and REV7 function in a common DNA replication fork protection pathway

Since both REV7 and PARP1 have reported roles in DNA replication fork protection,^22,61^ we asked whether either protein functions in the same pathway as RNF25 to protect stalled forks. We used siRNA to knock down PARP1 or REV7 in *RNF25^+/+^*or *RNF25^-/-^* H1299 cells (Supplementary Fig. 4b). Then we performed DNA fiber assays to measure fork movement and fork degradation in the resulting cells. Consistent with previous studies, loss of either PARP1 or REV7 resulted in a decrease in fork speed in *RNF25^+/+^*cells (Fig. 6a). Knockdown of PARP1 in *RNF25^-/-^* cells resulted in an additional decrease in fork speed, while knockdown of REV7 did not affect fork speed in cells lacking RNF25 (Fig. 6a). Similarly, in the fork degradation assay, knockdown of PARP1 or REV7 in *RNF25^+/+^* resulted in increased fork degradation (Fig. 6b). However, knockdown of PARP1 but not of REV7 in RNF25^-/-^ cells resulted in an additional increase in fork degradation (Fig. 6b). Thus RNF25-deficiency phenocopies the DNA replication defects of PARP1 or REV7-depleted cells. However, combining RNF25-deficiency with PARP1-depletion has additive effects on DNA replication stress parameters, while the effects of RNF25-deficiency in combination with REV7-depletion are non-additive. We conclude that RNF25 and PARP1 function in separate pathways, while RNF25 and REV7 are epistatic and function in a common pathway to promote DNA replication stress tolerance.

**Fig. 6:**
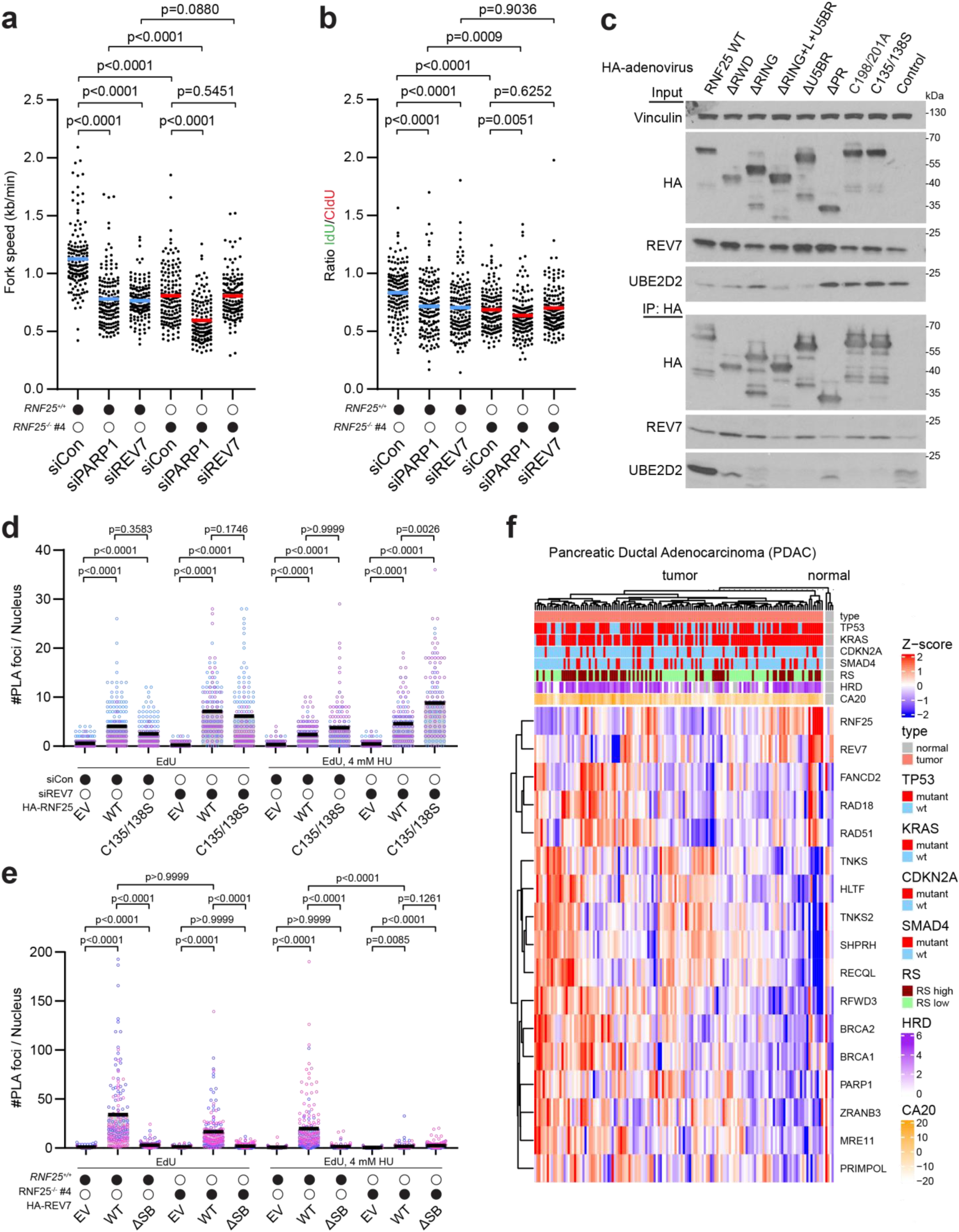
RNF25 and REV7 function in a common DNA replication fork protection pathway. **a, b** *RNF25^+/+^* and *RNF25^-/-^* H1299 cells were transfected with indicated siRNAs 48 h before conducting the DNA fork movement assay (a) and the fork degradation assay (b). 150 fibers were analyzed per condition with means indicated with bars. Each figure shown is a representative experiment from two biological replicates. Statistics: two-tailed Mann-Whitney test. **c** HA-RNF25 WT and domain mutants were expressed in H1299 cells using adenoviral vectors, then immunoprecipitated with anti-HA beads. **d** SIRF assay between HA-RNF25 (WT and C135/138S) and EdU in *RNF25^-/-^*H1299 cells transfected with control or REV7 siRNA. The number of foci per nucleus was quantified using ImageJ. Each biological replicate is shown in purple and blue, with means indicated using bars. Number of nuclei quantified per condition (from left to right) (replicate 1-purple, replicate 2-blue): (99, 64); (75, 96); (77, 75); (84, 74); (76, 82); (78, 61); (88, 86); (87, 80); (75, 70); (81, 66); (76, 77); (77, 63). Statistics: Kruskal-Wallis test with Dunn’s multiple comparisons test. **e** SIRF assay between HA-REV7 (WT and ΔSB) and EdU in *RNF25^+/+^* and *RNF25^-/-^* H1299 cells. The number of foci per nucleus was quantified using ImageJ. Each biological replicate is shown in blue and pink with means indicated using bars. Number of nuclei quantified per condition (from left to right) (replicate 1-blue, replicate 2-pink): (70, 75); (74, 77); (71, 87); (70, 76); (73, 85); (72, 76); (70, 76); (75, 86); (72, 83); (74, 75); (65, 80); (72, 79). Statistics: Kruskal-Wallis test with Dunn’s multiple comparisons test. **f** Heatmap depicting relative mRNA expression of replication and DNA damage repair factors in PDACs from TCGA. Samples were clustered as normal or tumor samples, with additional classification of tumor samples by mutation status of key PDAC driver genes and genome maintenance related features (RS – Replication Stress signature, HRD – Homologous Recombination Deficiency score, CA20 – Centrosome Amplification signature).

Next we performed co-immunoprecipitation experiments using a panel of HA-RNF25 mutants to identify the domains of RNF25 required for the interaction with REV7. As shown in Fig. 6c the association of REV7 with HA-RNF25 mutants lacking the RING+L+U5BR domains, or the Proline-Rich domain was reduced (by 2.33-fold and 3.13-fold respectively) when compared with wild-type HA-RNF25. Critically, the HA-RNF25 C135/138S mutant which lacks E2-binding activity retained REV7 binding (Fig. 6c). These results show good correlation between the REV7-binding activity of HA-RNF25 mutants and their ability to complement the DNA replication defects of *RNF25^-/-^* cells.

Based on our demonstration of an epistatic relationship, we next determined whether RNF25 and REV7 were interdependent for recruitment to sites of DNA synthesis. First we used SIRF assays to determine the effect of REV7 knockdown (Supplementary Fig. 4c) on localization of wild-type HA-RNF25 or HA-RNF25 C135/138S in relation to nascent DNA. As shown in Fig. 6d, both wild-type HA-RNF25 and the E2 interaction-deficient HA-RNF25 C135/138S mutant localized to newly replicated DNA under basal conditions and following replication stress. For both wild-type and C135/138S HA-RNF25, localization to nascent DNA was not inhibited by REV7-depletion (Fig. 6d). We conclude that REV7 is not required for the recruitment of RNF25 to DNA replication forks. Since REV7 has been shown to interact with newly replicated DNA and protect replication forks^22^ we also asked whether the recruitment of REV7 to nascent DNA is RNF25 dependent. We used the SIRF assay to measure the proximity between HA-tagged REV7 and EdU-labelled DNA in both *RNF25^+/+^* and *RNF25^-/-^* H1299 cells. Under basal conditions, we observed interaction between REV7 WT and nascent DNA in both *RNF25^+/+^* and *RNF25^-/-^* backgrounds. As a specificity control for REV7 localization, we showed that a REV7 ‘ΔSB’ mutant lacking the 10 C-terminal amino acids of the ‘seatbelt’ domain which mediates binding to its partners^67^ failed to localize to nascent DNA (Fig. 6e, Supplementary Fig. 4d). When cells were treated with HU following the EdU pulse, we observed significantly reduced PLA foci between REV7 and EdU in *RNF25^-/-^*cells when compared with *RNF25^+/+^* cells. These results suggest that the roles of REV7 in basal fork movement and in fork protection following replication stress are separable, and that REV7 depends on RNF25 to localize to DNA following replication stress.

To determine whether *RNF25* expression correlates with other fork reversal or fork protection factors in pathological settings, we analyzed gene expression datasets from The Cancer Genome Atlas (TCGA). In the three tumor types that we interrogated, Pancreatic Ductal Adenocarcinoma (PDAC) (Fig. 6f), Lung Adenocarcinoma (LUAD) and Breast Invasive Carcinoma (BRCA) (Supplementary Fig. 4f, g), *RNF25* and *REV7* expression were the most closely correlated, with a significant positive Pearson’s correlation coefficient (Supplementary Fig. 4h). The co-expression of *RNF25* and *REV7* in patient tumors is also consistent with our results showing that RNF25 and REV7 cooperate in the same pathway to promote DNA replication stress tolerance.

### RNF25 and Polζ function in a common DNA replication fork protection pathway

REV7 protects reversed DNA replication forks independently of Shieldin and in a REV1- and REV3L-dependent manner.^22^ Given our finding that REV7 and RNF25 function in the same fork protection pathway, we performed epistasis experiments to determine the genetic relationships between RNF25 and REV3L, REV1, and Shieldin. As shown in Fig. 7a, knockdown of SHLD2, REV1, and REV3L (Supplementary Fig. 4e) resulted in increased fork degradation in *RNF25^+/+^*cells. However, in *RNF25^-/-^* cells, knockdown of SHLD2 resulted in an additive increase in fork degradation, whereas knockdown of REV1 and REV3L had no additive effect (Fig. 7a). These results suggest that similar to REV7, RNF25 functions in the same pathway as REV1 and REV3L.

**Fig. 7:**
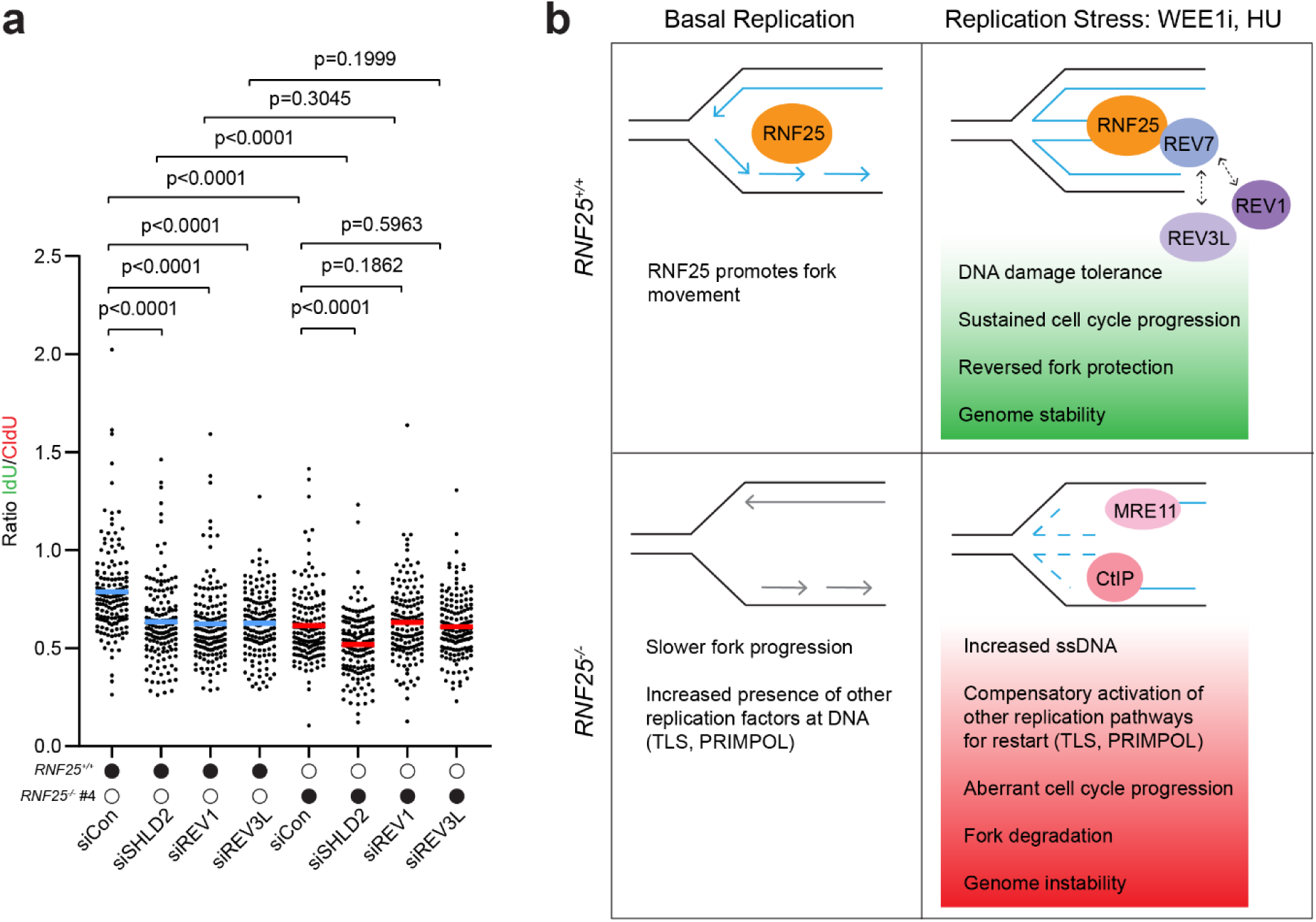
RNF25 and Polζ function in a common DNA replication fork protection pathway. **a** *RNF25^+/+^* and *RNF25^-/-^* H1299 cells were transfected with indicated siRNAs for 48 h before conducting the DNA fiber fork degradation assay. 150 fibers were analyzed per condition with means shown using bars. This figure is a representative experiment from two biological replicates. Statistics: two-tailed Mann-Whitney test. **b** Schematic depicting the role of RNF25 in basal replication and following replication stress. Under basal conditions, RNF25 promotes fork movement. Under replication stress conditions, RNF25 protects reversed forks by recruiting REV7. REV1 and REV3L also function in the same fork protection pathway. However, in RNF25 deficient cells, fork degradation is mediated by nucleases MRE11 and CtIP.

Taken together, our results identify RNF25 as a new mediator of DNA replication fork movement and replication fork protection (Fig. 7b). We propose a model in which RNF25 facilitates replication fork progression and that in its absence, basal replication movement is slowed and ssDNA accumulates. Following replication stress, RNF25 recruits REV7 to replication forks and together with REV1 and REV3L, protects reversed replication fork structures from degradation. In the absence of RNF25, there occurs extensive degradation of the replication fork by nucleases MRE11 and CtIP. Importantly, the E2-binding activity of RNF25 is dispensable for fork protection, thereby defining a non-canonical role that is separable from its known functions in protein ubiquitylation.

## Discussion

Our findings expand on the model of replication stress tolerance proposed by Paniagua et al.^22^ in that we identify RNF25 as a binding partner and critical upstream regulator of REV7-mediated fork protection. Similar to the reported role of REV7 in fork protection,^22^ we show that RNF25 is epistatic to REV1 and REV3L, but not the Shieldin complex.

Paniagua et al.^22^ found that the catalytic activity of REV3L was required for fork protection and proposed that REV7 and REV3L might either physically block the access of nucleases to reversed forks or promote resynthesis of resected DNA. The resynthesis model may be supported by the accumulation of ssDNA after loss of REV7 or REV3L.^22^ We also observed aberrantly high levels of ssDNA and pRPA in *RNF25^-/-^* cells experiencing replication stress (Fig. 2). However, the exact source of ssDNA in RNF25-deficient or REV7-deficient cells is not yet clear and needs further investigation. Another possibility is that RNF25 functions upstream of REV7/REV3L (which constitute DNA polymerase ζ) and REV1 to promote sealing of ssDNA gaps that remain after replication fork restart. The latter hypothetical mechanism may be consistent with the canonical TLS functions of REV1 and Polζ in ssDNA gap filling.

We show that the E2 ligase-binding activity of RNF25 is dispensable for its function in fork protection. In addition to their roles in tethering E2 ubiquitin ligases to their substrates^65,68^ some RING finger E3 ligases participate in DNA repair via molecular chaperone and scaffold activities, independently of stimulating ubiquitin conjugation. For example, RAD18 plays several roles as a molecular chaperone for RAD51 in homologous recombination, SLF2 in interstrand crosslink repair, and Pol η in TLS.^69–71^ Additionally, RNF8 recruits CHD4 (the catalytic subunit of the NuRD complex) to damaged chromatin through a non-canonical interaction to promote chromatin decondensation and accessibility of other repair factors.^72^ Even in their canonical roles in promoting ubiquitylation, RING-finger E3 ligases lack any intrinsic catalytic activity and instead serve as scaffolds that physically link E2 ubiquitin-conjugating enzymes to their substrates.^65,68^ It is likely that RING-finger E3 ligases have broader roles as molecular scaffolds than are currently appreciated.

Replication fork reversal is a protective mechanism that stabilizes the replication fork during stalling. However, the reversed fork structure contains free DNA ends that are potential substrates for many nucleases.^17^ We show here that RNF25 protects reversed forks against degradation by a specific subset of nucleases which includes CtIP and MRE11. Our results are consistent with previous findings that different fork remodelers generate reversed fork structures that are degraded by distinct nucleases and guarded by different fork protection factors.^21^ For example, fork remodelers HLTF, SMARCAL, and ZRANB3 operate in a separate pathway from FBH1, and each pathway generates DNA structures that are safeguarded by distinct fork protectors. Thus, 53BP1 protects FBH1-dependent reversed forks from degradation by DNA2 whereas BRCA2 and FANCD2 protects reversed forks generated by SMARCAL1, ZRANB3, and HLTF from MRE11-dependent degradation.^21^

In our experiments, depleting HLTF or MRE11 rescued the fork degradation phenotype of *RNF25^-/-^* cells (Fig. 4c). Our finding that CtIP-depletion rescues the fork degradation phenotype of RNF25^-/-^ cells may be explained by previous work showing that CtIP promotes MRE11-dependent degradation of reversed forks.^15,73^ Taken together our results suggest that RNF25 protects HLTF-induced regressed forks from nucleolytic attack by MRE11. HLTF is dispensable for fork degradation when the fork protection factor 53BP1 is inactivated.^21^ Moreover, knockdown of FBH1 or DNA2 did not rescue the fork degradation phenotype of REV7-deficient cells.^22^ Therefore, we consider it unlikely that RNF25 protects regressed forks generated through the FBH1-dependent remodeling pathway.

We observed increased levels of chromatin-bound MUS81 in *RNF25^-/-^* cells, both basally and after HU-induced replication stress (Fig. 4a, b). However, knockdown of MUS81 had no effect on fork degradation in *RNF25^-/-^*cells (Fig. 4c). It is possible that the increased chromatin loading of MUS81 in *RNF25^-/-^*cells is related to the restart of reversed forks rather than fork degradation. Similarly, in *BRCA2* deficient cells, MUS81 loss did not affect fork degradation. However, partially resected forks generated by MRE11 were subsequently cleaved by MUS81 to promote POLD3 dependent fork restart in BRCA2 deficient cells.^15^ Further studies are needed to determine if reversed forks are restarted via a MUS81 and POLD3 dependent mechanism when RNF25 is absent.

Two other CRISPR screens have identified roles for RNF25 in tolerating genotoxic agents including MMS and UV.^74,75^ However, the role of RNF25 in tolerating those genotoxins may be related to UV- and MMS-induced mRNA damage and ribosome collisions. Recent work shows that RNF25, and another E3 ligase RNF14, are activated by the translational stress sensor GCN1 to remediate ribosome stalling.^51,76^ In response to ribosome collisions, RNF25 ubiquitinates the ribosomal protein RPS27A. Then, RNF14 ubiquitinates the elongation factor eEF1A, leading to proteosome-dependent degradation of eEF1A which clears the occluded ribosome ‘A site’.^51^ RNF25 and RNF14 also cooperate to target the termination factor eRF1 for degradation when stalled.

We consider it unlikely that the role of RNF25 at DNA replication forks is related to its role in translation quality control for several reasons. First, in our CRISPR screen, we used a WEE1 inhibitor as a source of replication stress. Mechanistically, WEE1 inhibition induces DNA damage in S-phase by derepressing CDK2 activity, leading to excessive origin firing.^77,78^ To our knowledge WEE1-inhibition is not known to induce RNA damage or ribosome collisions. Second, we show that RNF14 is dispensable for tolerating DNA replication stress (Supplementary Fig. 2e). Finally, the role of RNF25 in translation quality control is dependent on its association with E2 ubiquitin conjugases, whereas E2-binding is dispensable for RNF25-dependent fork protection. Nevertheless, the participation of RNF25 in both DNA replication and translation quality control may represent a mechanism for integrating these important cellular processes in response to stress.

We speculate that the RNF25-PARP interaction we identified is relevant for RNF25 functions in translation. Although PARP1 has several known roles in DNA replication,^57–61^ our epistasis experiments suggest that RNF25 and PARP1 do not act in the same pathway for fork movement or fork protection (Fig. 6a, b). However, PARP enzymes have key roles in ribosome biogenesis, mRNA processing, and translation.^79–82^ Therefore, it will be interesting to investigate the possible role of RNF25-PARP1 signaling in ribosome function.

Given that RNF25 mediates processes that are critical for proliferation and stress adaptation of cancer cells (genome maintenance and translation quality control) it is interesting to consider RNF25 and its distal effectors as therapeutic target pathways. E3 ligases are emerging as attractive therapeutic targets^30–33^ and it will be important to determine whether RNF25 and its effectors are pharmacologically tractable. WEE1 inhibitors are being used as experimental therapies for several cancers.^37,38,83,84^ Nucleotide depletion (modeled by our HU treatment experiments) is a common feature of oncogene-driven cancers and a wide range of nucleotide synthesis inhibitors are being used in cancer therapies.^7,8^ RNF25 inhibition could also be an attractive combination therapy for use with existing pharmacological stressors.

## Methods

### Cell culture and transfection

Pa02C and Pa03C cells were kind gifts from Dr. Channing Der (UNC Chapel Hill). H1299 cells were purchased from the American Type Culture Collection (ATCC). Pa02C, Pa03C, and H1299 cells were cultured in Dulbecco’s Modified Eagle’s Medium (DMEM, Corning 10-017-CV) supplemented with 10% fetal bovine serum (Gibco 10437-028), 100 U/ml penicillin, and 100 µg/ml streptomycin (Gibco 15-140-122) at 37°C, 5% CO_2_. Plasmid and siRNA transfections were done using Lipofectamine 2000 (Invitrogen 11668019) according to manufacturer’s protocol, except for plasmid transfections where amounts of lipofectamine and plasmid were halved to reduce toxicity.

siRNAs used in this study are listed with UU overhangs included: siControl: UAGCGACUAAACACAUCAAUU; siRNF25-A: GAGGGAGGCAAUAAAGAUAUU, siRNF25-D: GGAACAGGGAAUUGGGGAUUU; siPARP1: AAGAUAGAGCGUGAAGGCGAAUU, siREV7 CAACACUGUCUGUCUCAAAUAUU. The following siRNAs were purchased from Dharmacon as SMARTpools: siMRE11: M-009271-01-0050; siMUS81: L-016143-01-0005; siCtIP: M-011376-00-0050, siEXO1: M-013120-00-0020; siHLTF: L-006448-00-0005; siSHLD2: L-013761-00-0005; siREV1: L-008234-00-0020; siREV3L: L-006302-00-0005; siSTAT1: L-003543-00-0005.

### Expression plasmids

The C-terminal Flag-HA-tagged REV7 WT and REV7ΔSB in the pCDH plasmid used for transient transfection were kind gifts from Dr. Junya Tomida (UNC Charlotte). The REV7ΔSB mutant has a 10 amino acid deletion from the C-terminus. All other lentiviral, adenoviral, and luciferase reporter plasmids are described in their respective sections.

### Adenovirus generation

N-terminal HA-tagged RNF25 wildtype and mutants (described in Fig. 5a) were cloned into the pACCMV vector^85^ and co-transfected in 239T cells with the pJM17 adenovirus plasmid.^85^ Adenovirus was precipitated from 293T cell lysates using polyethylene glycol. Adenovirus was further purified by CsCl gradient centrifugation and gel filtration chromatography. Purified adenovirus was added directly to the cell culture medium for cell infection.

### Doxycycline-inducible stable expression of RNF25 domain deletions and mutants

N-terminal HA-tagged RNF25 wildtype and mutants (described in Fig. 5a) were cloned into the pINDUCER20 plasmid^86^ and co-transfected with packaging plasmid ΔNRF and envelope plasmid pMDK64 in HEK293T cells to produce lentivirus. *RNF25^+/+^*and *RNF25^-/-^* H1299 cells were infected with lentivirus along with 8 μg/ml polybrene (Sigma Aldrich TR-1003-G) for 24 h before plating at low density in media with 400 μg/ml G418 for selection. Selected pools were used for downstream assays.

### E3 Ubiquitin Ligase CRISPR-Cas9 loss-of-function screen library design

The E3 ubiquitin ligase CRISPR-Cas9 library contains 3,122 sgRNAs targeting 317 RING-domain containing genes, several SUMO ligases and genes of interest, and 1,000 non-targeting sgRNAs. Each gene has ten domain-focused sgRNAs and each oligo (74 nt) contains 20 nt target sgRNA, 5’ universal flanking sequence (GTGGAAAGGACGAAACACCG), and a 3’ universal flanking sequence (GTTTTAGAGCTAGAAATAGCAAGTTAAAATAAGG).

### E3 ubiquitin ligase CRISPR library cloning

E3 library sgRNA pool was cloned into lentiCRISPR v2 (Addgene #52961)^87^ via NEBuilder HiFi DNA Assembly. PCR amplification of the library sgRNA pool were performed using 017_ArrayF (TAACTTGAAAGTATTTCGATTTCTTGGCTTTATATATCTTGTGGAAAGGACGAAACACCG) and 018_ArrayR (ACTTTTTCAAGTTGATAACGGACTAGCCTTATTTTAACTTGCTATTTCTAGCTCTAAAAC) primers. PCR protocol: Initial denaturation (98°C, 30 s), Denaturation (98°C, 10 s), Annealing (65°C, 30 s), Extension (72°C, 25 s), Final Extension (72°C, 5 min). Denaturation, annealing, and extension steps were cycled 30 times. 50 µL PCR reaction: 2.5 µL of each primer (10 µM stock), 2 µL E3 library pool (0.2 µM stock), 18 µL molecular biology grade water, 25 µL 2X Q5 High-Fidelity 2X Master Mix (NEB, M0492L). PCR products were purified using the MinElute PCR purification kit (QIAGEN, 28004). Plasmid digestion: lentiCRISPR v2 vector was digested with BsmBI (NEB, R0580) at 55°C overnight then heat inactivated at 80°C for 20 min. Digestion products were run on a 0.8% agarose gel and a 13 kb fragment was extracted using a gel extraction kit (QIAGEN, 28706). HiFi assembly reaction: 100 ng BsmBI digested vector (2 µL), 6.66 ng E3 library pool (8 µL) and 10 µL HiFi DNA Assembly Master Mix (NEB, E2621). Reaction mixture was incubated at 50°C for 1 hour. Electroporation: 0.75 µL of HiFi assembly mixture was added to 25 µL of electrocompetent bacteria (Lucigen, 60242-2). Bacteria and DNA mixture was electroporated in ice-chilled cuvettes (Bio-Rad, 1652083) using Gene Pulser Xcell electroporator (Bio-Rad, 1652660) at 1800 Volts, 10 µFarad, 600 Ohm, 1 mm cuvette gap. 500 µL of recovery media was added immediately post electroporation (Lucigen, 80026-1). 6 electroporation reactions were performed to ensure high coverage of the entire library. Transformed bacteria were incubated at 37°C for 1 hour then plated on 12 LB ampicillin 15 cm plates to incubate overnight at 37°C. 10 ml of LB was added to each plate and transformed bacteria were removed using sterile scrapers. Bacteria from 3 plates were transferred to 500 ml LB cultures and incubated at 37°C for 3 hours. Cloned plasmid library was extracted using a plasmid maxiprep kit (QIAGEN, 12362).

### CRISPR-Cas9 screening

Pa02C and Pa03C were seeded in 6-well plates and transduced with the E3 ubiquitin CRISPR-Cas9 library at a multiplicity of infection (MOI) of 0.3 with 300x sgRNA coverage (300 cells per sgRNA). Transduced cells were selected by puromycin (Gibco A1113803) for 5-6 days after reseeding in 15 cm dishes. Selected cells were treated with DMSO or MK-1775 at IC_30_ for 20 population doublings (PD) as measured by cell counting (Thermo Fisher Scientific, Countess II). Cell pellets were collected at PD10 and PD20 and stored at -80°C.

### Indexed sgRNA pool preparation and sequencing

Genomic DNA (gDNA) was isolated from cell pellets (QIAGEN, 69506). Two PCR reactions were performed in succession to (1) amplify sgRNAs from isolated gDNA and (2) index sgRNA pools from each sample. The sgRNA amplification PCR was performed using 8 µg of gDNA and an equimolar forward primer mixture (CGACGCTCTTCCGATCTCGATTTCTTGGCTTTATATATCTTGTGGAAAGG, CGACGCTCTTCCGATCTTCGATTTCTTGGCTTTATATATCTTGTGGAAAGG, CGACGCTCTTCCGATCTATCGATTTCTTGGCTTTATATATCTTGTGGAAAGG and CGACGCTCTTCCGATCTGATCGATTTCTTGGCTTTATATATCTTGTGGAAAGG) and a reverse primer (CGTGTGCTCTTCCGATCCAATTCCCACTCCTTTTCAAGACCTAG) using 2X Q5 High-Fidelity 2X Master Mix (NEB, M0492L). sgRNA amplification PCR protocol (50 µL): Initial denaturation (98°C, 30 s), Denaturation (98°C, 15 s), Annealing (61°C, 30 s), Extension (72°C, 30 s), Repeated 9 times, Denaturation (98°C, 15 s), Annealing (68°C, 30 s), Extension (72°C, 30 s), Repeated 19 times, Final Extension (72°C, 5 min). Indexing PCR was performed using 1 ng of sgRNA pool mixture and Illumina dual index forward and reverse primers with adapters (i7 forward primer, CAAGCAGAAGACGGCATACGAGAT[i7-seq]GTGACTGGAGTTCAGACGTGTGCTCTTCCGATC and i5 reverse primer, AATGATACGGCGACCACCGAGATCTACAC[i5-seq]ACACTCTTTCCCTACACGACGCTCTTCCGATCT) using 2X Q5 High-Fidelity 2X Master Mix (NEB, M0492L). Indexing PCR protocol (100 µL): Initial denaturation (98°C, 30 s), Denaturation (98°C, 15 s), Annealing (60°C, 30 s), Extension (72°C, 45 s), Repeated 14 times, Final Extension (72°C, 5 min). After each PCR reaction, PCR products were size selected using Ampure XP bead mixture (Beckman Coulter, A63881) according to manufacturer’s instructions with a 1:2 and 1:1 bead mixture to PCR product ratios for sgRNA amplification and indexing PCR, respectively.

### CRISPR-Cas9 screen analysis

Multiplexed next-generation sequences were demultiplexed into individual FASTQ sample files. The sgRNA count and statistical significance was analyzed using a previously published algorithm -Völundr v2.1.0.^47^ Völundr-formatted files for the i5 and i7 dual indexes, sample indexes, E3 ligase sgRNA sequences, sgRNA sequence target search and statistic settings can be found on Zenodo and are publicly available as of the date of publication at (10.5281/zenodo.13930006). sgRNA abundance scores and p-values were calculated using the plasmid library as a control.

### Target gene knock-out cell line generation

To generate non-targeting control, *TRIM11^-/-^*, *RNF25^-/-^* and *PIAS1^-/-^* PDAC polyclonal cell lines, Pa03C and Pa02C cells were infected with lentivirus produced in HEK293T cells (from the lentiCRISPR v2 plasmid encoding sgRNAs and Cas9) with polybrene (Sigma Aldrich TR-1003-G) pre-treatment on cell monolayers at 8ug/ml for 30 minutes. Stably transduced cells were split after 24 h and selected in 1 μg/ml puromycin for 7 days. To generate *RNF25^-/-^* clones in H1299 cells, cells were transfected with an equal mix of the 3 pLENTI CRISPR plasmids with guides targeting RNF25 for 6 hours. Cells were then selected in puromycin (2 μg/ml) for two days. Selection was stopped when the plasmid was lost from cells, tracked by a parallel GFP transfection. Single clones were generated by plating cells in low density in 96-well plates (1 cell/well). After clones were expanded, they were verified to be puromycin sensitive, to ensure that the Cas9 gene was not stably integrated so that rescue experiments could be performed in these cells.

sgRNA oligos were purchased from IDT and Eton Bioscience.

NT sg1 GTGGAAAGGACGAAACACCGCTTTTCAGCTGAGACGTACGGTTTTAGAGCTAGAAATAGCAAGTTAAAATAAGG

NTsg3 GTGGAAAGGACGAAACACCGCGGAGTTAACCTGGAACCTTGTTTTAGAGCTAGAAATAGCAAGTTAAAATAAGG

NT sg4 GTGGAAAGGACGAAACACCGAATCGACTCGAACTTCGTGTGTTTTAGAGCTAGAAATAGCAAGTTAAAATAAGG

RNF25 sg1 GTGGAAAGGACGAAACACCGATCTCTATCCGAAATCCCCGGTTTTAGAGCTAGAAATAGCAAGTTAAAATAAGG

RNF25 sg3 GTGGAAAGGACGAAACACCGGGATTTCGGATAGAGATCTGGTTTTAGAGCTAGAAATAGCAAGTTAAAATAAGG

RNF25 sg7 GTGGAAAGGACGAAACACCGGGATGTACCGAGCAAGGCAGGTTTTAGAGCTAGAAATAGCAAGTTAAAATAAGG

PIAS1 sg1 GTGGAAAGGACGAAACACCGTACGCCGGGAGAAACAAGCAGTTTTAGAGCTAGAAATAGCAAGTTAAAATAAGG

PIAS1 sg2 GTGGAAAGGACGAAACACCGGAACATGTAAGGGCCCGACAGTTTTAGAGCTAGAAATAGCAAGTTAAAATAAGG

TRIM11 sg1 GTGGAAAGGACGAAACACCGGTTGCTGTTCCAAGCCCAGGGTTTTAGAGCTAGAAATAGCAAGTTAAAATAAGG

TRIM11 sg2 GTGGAAAGGACGAAACACCGCGGGACTGGTAGAGACACTGGTTTTAGAGCTAGAAATAGCAAGTTAAAATAAGG

### Colony formation assay

Cells were seeded at a low density of 2000 cells/well in triplicate in six-well dishes. Drugs were diluted in growth medium and added directly to the cells after 24 h of seeding and replenished every 3 days during media changes (WEE1i: MedChemExpress HY-10993). All dishes were harvested when the control group was about 80% confluent, typically between 10 to 14 days after seeding. Cells were stained with 0.05% crystal violet in 1x PBS containing 1% methanol and 1% formaldehyde. Fiji plugin ColonyArea^88^ was used to automatically quantify the scanned images of stained colonies at a constant threshold for respective experiments.

### Immunoblotting

To prepare extracts containing soluble and crude chromatin-associated proteins, monolayers of cultured cells were washed in ice-cold PBS and lysed in ice-cold cytoskeleton buffer (CSK buffer: 10 mM Pipes, pH 6.8, 100 mM NaCl, 300 mM sucrose, 3 mM MgCl_2_, 1 mM EGTA, 25 mM β-glycerophosphate, 10 mM NaF, and 0.1% Triton X-100) freshly supplemented with protease inhibitor cocktail (Roche 4693159001) and PhosSTOP (Roche 4906837001). Lysates were centrifuged at 4,000 rpm for 8 min to separate the soluble fraction from CSK-insoluble nuclei. The detergent-insoluble nuclear fractions were then resuspended in a minimal volume of CSK. Both soluble and insoluble lysate samples were normalized in Laemelli buffer based on protein measurements from a Bradford assay and heated to 95°C for 8-10 min. Chromatin samples presented in Fig. 4a and 4b are proteins released from the chromatin after nuclease digestion. The crude chromatin fraction was washed twice with CSK, then incubated with 250U/ml nuclease (Pierce 88702) for 30 min on ice. Samples were then spun at 15,000g for 10 min to obtain proteins released from chromatin digestion.

Protein samples were then separated by SDS-PAGE, transferred to nitrocellulose membranes, blocked for 1 h in 5% nonfat dried milk diluted in TBST followed by overnight incubation with indicated primary antibodies. Antibodies used: TRIM11 (ab111694), pIRF3 S386 (ab76493), MUS81 (ab14387), and MRE11 (ab214) from Abcam; RNF25 (A303-844A), pRPA S33 (A300-246A), pRPA S4/S8 (A300-245A), ZRANB3 (A303-033A), HLTF (A300-230A), Pol η (A301-231A), RAD18 (A301-340A), and HA (A190-138A) from Bethyl Laboratories; Vinculin (V4505) and γH2AX (05-636) from Sigma Millipore; RPA 34 (NA19L) from Calbiochem; pATM S1981 (sc-47739), β-actin (sc-47778), GAPDH (sc-32233), TNKS1/2 (sc-365897), PARP1 (sc-8007), UBE2D2 (sc-100617), REV1 (sc-393022), FANCD2 (sc-28194), and PCNA (sc-56) from Santa Cruz Biotech; CDC2 pTyr15 (CST 9111), PIAS1 (CST 3550), Phospho-p44/42 MAPK (Erk1/2) (Thr202/Tyr204) (CST 9106), pSTING Ser366 (CST 19781), STING (CST 13647), pSTAT1 Tyr701 (CST 9167) and STAT1 (CST 9175) from Cell Signaling Technology; REV7 (12683-1-AP), EXO1 (16253-1-AP), and PRIMPOL (29824-1-AP) from Proteintech; CtIP (61141) from Active Motif. Next morning, the blots were washed 3x in TBST for 10min/wash, incubated in secondary anti-rabbit (Bethyl A120-101P), anti-mouse (Bethyl A90-116P), or anti-goat (Santa Cruz Biotech sc-2020) HRP antibody for 1-2 h, and washed 3x in TBST. Western Lightning™ Plus Chemiluminescence Reagent (Revvity NEL104001EA) was added to membranes for 1-2 min and proteins visualized with a Li-Cor Odyssey Imager or using autoradiography film.

### RT-qPCR

Total RNA was isolated using a RNeasy Mini Kit (Qiagen 74106), cDNA was synthesized using the iScript cDNA Synthesis Kit (Bio-Rad 1708891), and qPCR performed using the iTaq Universal SYBR Green Supermix (Bio-RAD 172-5121) and the ABI 7500 Fast Real-Time PCR System according to manufacturer’s instructions. The following primers were used: REV3L: forward TCATGAGAAGGAAAGACACTTTATG, reverse GCTGTAGGAGGTAGGGAATATG; SHLD2: forward ACTGACAGAGGCAGTATACAGTT, reverse AGGAAATGCCAGCTCTGAAA; GAPDH: forward GTCTCCTCTGACTTCAACAGCG, reverse ACCACCCTGTTGCTGTAGCCAA. Results were analyzed using the 2-ΔΔCt method, normalized to GAPDH.

### Cell cycle distribution assay

*RNF25^-/-^* and *PIAS1^-/-^* Pa03C polyclonal cell lines, along with *RNF25^+/+^* and *PIAS1^+/+^* Pa03C control cells, were treated with DMSO or WEE1i for 24 h. Cells were collected by trypsinization, fixed in 35% ethanol overnight at 4°C. The lysates were then washed twice with DPBS and stained with 1mg/ml propidium iodide (Invitrogen #P3566) with 8ug/ml RNaseA (Sigma Aldrich R4642) to stain DNA. Unstained samples were used to distinguish cell populations. Stained nuclei were assayed by flow cytometry on a Thermo Fisher Attune NxT analyzer and data was analyzed in Flowjo v10.8 software.

### ssDNA assay

To detect ssDNA, sub-confluent cells were cultured for 36 h in medium containing 10 μM BrdU to label genomic DNA, in duplicate. Following BrdU labeling, media was changed to fresh BrdU-free medium and cells were incubated overnight for 12 h. WEE1i was diluted in fresh media and incubated with cells for another 24 h. Then cells were harvested by trypsinization, resuspended in 65% PBS with 35% ethanol, and fixed overnight at 4°C. Fixed cells were washed with PBS, resuspended in blocking buffer (0.25% Tween20 and 0.5% BSA in PBS) and incubated with primary pH3 antibody (Upstate/Millipore 06-570) (1:200 in blocking buffer) for 2 h rotating at 4°C. The samples were then washed with PBS followed by blocking buffer. Subsequently, samples were incubated with 4:10 FITC conjugated anti-BrdU antibody (BD Biosciences 556028) and 1:200 Alexa anti-Rabbit 647 (Life technologies A31573) secondary antibody in blocking buffer for 1 h at RT to detect pH3 antibody. The samples were then incubated overnight in PBS containing 10 μg/ml of propidium iodide (Invitrogen P3566) and 8 μg/ml of RNaseA (Sigma Aldrich R4642) to stain total DNA. For FACS controls, unstained sample, FITC-BrdU antibody only sample without denaturation, FITC-BrdU only sample with denaturation by HCl (followed by neutralization with Borax and PBS wash) and pH3 antibody-Alexa 647 only sample were prepared. Stained cells were run in a Thermo Fisher Attune NxT analyzer and data was analyzed in FlowJo v10.8 software.

### Live cell imaging

*RNF25^-/-^* and *PIAS1^-/-^* Pa03C polyclonal cell lines and *RNF25^+/+^* and *PIAS1^+/+^*Pa03C control cells stably expressing H2B-GFP (Addgene #69550) were generated with polybrene assisted transduction and infected cells were selected by flow sorting with the BD FACSAria II. Cells were seeded on Chambered Coverglass (Nunc 155382) or 24-well glass bottom plates (Greiner 662892) for live cell imaging. 12 h later cell monolayers were incubated in fresh media with WEE1i for 36 h. Time-lapse microscopy was performed on a Keyence BZ-X810 using a 20x objective. Images were taken at 3 min intervals for 24 h. Best focus projections of the time series were exported into AVI format. Image sequences were generated using ImageJ and cell fates manually tracked and quantified.

### DNA fiber assays

Cells were transfected with siRNA for 48 h before being pulsed-labeled for DNA fiber assays. Cells with doxycycline-inducible genes were cultured in media with 100 ng/ml doxycycline for 48 h prior to pulse labeling. For the fork movement assay, exponentially growing cells were pulsed with 25 μM CldU (MedChemExpress HY-112669) for 20 min, washed 2x with media containing 250 μM IdU (MedChemExpress HY-B0307), then pulsed with 250 μM IdU for 20 min. For the fork degradation assay, cells were pulsed with CldU followed by IdU for 30 min each, then treated with 4 mM HU (Sigma-Aldrich 400046) for 4 h. Cells were then washed 2x with PBS, trypsinized and pelleted at 1600rpm for 4 min, and resuspended in ice-cold PBS at a concentration of 2.5×10^5^ cells/ml. 2 μl of cells were pipetted to a glass slide and left to air dry for about 5 minutes. Then 9 μl of spreading buffer (200 mM Tris pH 7.4, 50 mM EDTA, 0.5% SDS) was added to the cell droplet and incubated for 2 min. Slides were tilted to spread the DNA at a low and constant speed. Slides were air dried for 2 min, then fixed in a 3:1 mix of methanol and acetic acid for 10 min and air dried for 10 min. Slides were stored at 4°C until staining.

Fixed slides were washed 2x with water, then denatured with 2.5 M HCl for 1 h 15 min. Slides were then rinsed 3x with PBS and incubated in blocking solution (1% w/v BSA in PBS-T (0.1% v/v Tween-20)) for 1 h. Slides were incubated with primary antibodies (1:500 rat anti-BrdU, Abcam ab6326; 1:600 mouse anti-BrdU, BD Biosciences 347580) for 1 h at room temperature before being washed 3x with PBST. Slides were then incubated with secondary antibodies (1:500 anti-rat Alexa Fluor 555, Invitrogen A21434; 1:600 anti-mouse Alexa Fluor 488, Invitrogen A11029) for 1.5 h at room temperature before being washed 3x with PBST, 2x with PBS, and mounted using Fluoroshield (Sigma-Aldrich F6182). Slides were stored at -20°C until imaging. Images were acquired using an inverted spinning disk confocal (Andor Dragonfly/Leica) with a 100x oil objective. Lengths of fibers were measured manually using the freehand line tool on ImageJ. At least 150-200 fibers were scored for each experiment, and only fibers with clearly defined beginnings and ends were measured. For all DNA fiber experiments in this paper, only contiguous CldU-IdU tracts were measured.

### Immunoprecipitation-Mass Spectrometry

H1299 cells were infected with control adenovirus or adenovirus expressing N-terminal HA-tagged RNF25 for 48 h, then cell lysates collected for immunoprecipitation with anti-HA beads (Roche 11815016001). Three technical replicates were collected and analyzed per sample. Immunoprecipitated protein complexes were subjected to on-bead trypsin digestion, as previously described.^89^ After the last wash buffer step, 50 µl of 50 mM ammonium bicarbonate (pH 8) containing 1 µg trypsin (Promega) was added to beads overnight at 37°C with shaking. The next day, 500 ng of trypsin was added then incubated for an additional 3 h at 37°C with shaking. Supernatants from pelleted beads were transferred, then beads were washed twice with 50 µl LC/MS grade water. These rinses were combined with original supernatant, then acidified to 2% formic acid. Peptides were desalted with peptide desalting spin columns (Thermo) and dried via vacuum centrifugation. Peptide samples were stored at -80°C until further analysis.

#### LC/MS/MS Analysis

Each sample was analyzed by LC-MS/MS using an Easy nLC 1200 coupled to a QExactive HF (Thermo Scientific). Samples were injected onto an Easy Spray PepMap C18 column (75 μm id × 25 cm, 2 μm particle size) (Thermo Scientific) and separated over a 120 min method. The gradient for separation consisted of a step gradient from 5 to 36 to 48% mobile phase B at a 250 nl/min flow rate, where mobile phase A was 0.1% formic acid in water and mobile phase B consisted of 0.1% formic acid in ACN. The QExactive HF was operated in data-dependent mode where the 15 most intense precursors were selected for subsequent HCD fragmentation. Resolution for the precursor scan (m/z 350–1700) was set to 60,000 with a target value of 3 × 10^6^ ions, 100ms inject time. MS/MS scans resolution was set to 15,000 with a target value of 1 × 10^5^ ions, 75ms inject time. The normalized collision energy was set to 27% for HCD, with an isolation window of 1.6 m/z. Peptide match was set to preferred, and precursors with unknown charge or a charge state of 1 and ≥ 8 were excluded.

#### Data Analysis

Raw data were processed using the MaxQuant software suite (version 1.6.12.0) for peptide/protein identification and label-free quantitation.^90^ Data were searched against a Uniprot Reviewed Human database (downloaded October 2020, containing 20,381 sequences) using the integrated Andromeda search engine. A maximum of two missed tryptic cleavages were allowed. The variable modifications specified were N-terminal acetylation, oxidation of Met, ubiquitylation of Lys and phosphorylation of Ser/Thr/Tyr. Label-free quantitation (LFQ) was enabled. Results were filtered to 1% FDR at the unique peptide level and grouped into proteins within MaxQuant. Match between runs was enabled. Data filtering and statistical analysis was performed in Perseus software (version 1.6.14.0). Proteins with log_2_ fold change ≥ 1 and a p-value < 0.05 are considered significant.

### Co-immunoprecipitations

H1299 cells were infected with HA-RNF25 or control adenovirus at a concentration of 5×10^9^ pfu/ml (Fig. 3f) or 1×10^10^ pfu/ml (Fig. 6c) for 48 h. Whole cell lysates were obtained by scraping cells with ice-cold CSK buffer with protease and phosphatase inhibitors, supplemented with nuclease (as described in Immunoblotting). Cell lysates were normalized for protein concentration using the Bradford assay. Lysates were incubated with anti-HA magnetic beads (Pierce 88836, MBL M180-11) for 3 h at 4°C on rotating racks. Beads were washed 3x with CSK, rotating 5 min per wash at 4°C. Washed beads were resuspended in 2x Laemmli buffer and incubated for 5 min at 95°C to release and denature proteins for SDS-PAGE.

### Immunofluorescence

*RNF25^-/-^* H1299 cells growing on coverslips were infected with 1×10^9^ pfu/ml adenovirus expressing HA-tagged RNF25 domain deletions and mutations for 24 h before soluble extraction and fixation (20 mM PIPES pH 6.8, 10 mM EGTA, 0.2% Triton X-100, 1 mM MgCl_2_, 4% paraformaldehyde) for 20 min at room temperature. Cells were washed 2x with PBST (0.5% Triton X-100) and blocked with 3% BSA in PBST for 1 h. Cells were incubated in HA primary antibody (1:100 Bethyl A190-138A) diluted in 1% BSA PBST for 1 h 15 min, then washed 4x with PBST and incubated with secondary antibody (1:300 Invitrogen A32816) in 1% BSA PBST for 1h. Cells were washed 4x with PBST and mounted on slides with Fluoroshield (Sigma-Aldrich F6182) with 1 μg/ml DAPI and imaged with an inverted spinning disk confocal (Andor Dragonfly/Leica) with a 100x oil objective.

### SIRF assay

Cells were transfected with siRNA or ectopically expressed plasmids for 72 hours before pulsing with EdU. Cells with doxycycline-inducible genes were cultured in media with 100 ng/ml doxycycline for 48 h prior to pulse labeling. All cells were grown on coverslips to sub-confluent density before being pulsed with 10 uM EdU for 10 min. For cells also treated with HU, coverslips were washed 2x with PBS before fresh media with 4 mM HU (Sigma-Aldrich 400046) was added for 4 h. Ice cold pre-extraction buffer (20mM NaCl, 3mM MgCl_2_, 300mM Sucrose, 10mM PIPES, 0.5% Triton X-100) was added to cells for 5 min on ice, then cells were washed once with PBS and fixed with 4% PFA for 10 min at room temperature. Fixed cells were then washed 3x with ice cold PBS.

Fixed cells were washed once with 3% BSA in PBS before the click reaction (Invitrogen C10640) was done according to manufacturer’s protocol, except that the picolyl azide used was a 5:1 mixture of biotin-picolyl azide (Vector Laboratories CCT11675, 20 uM final concentration) and the provided Alexa Fluor 647 picolyl azide. After coverslips were incubated with the click reaction cocktail for 30 minutes at room temperature, cells were washed 3x with PBS and blocked with 5% BSA in PBS for 2 h at room temperature. Cells were incubated with primary antibodies HA (1:200 CST 3724S) and biotin (1:200 sc-101339) diluted in 10% FBS in PBS overnight at 4°C.

Next day, coverslips were incubated with Duolink PLA Probes (Sigma-Aldrich DUO92004 Mouse Minus, DUO92002 Rabbit Plus) and ligation and amplification (Sigma Aldrich DUO92014) carried out according to manufacturer’s instructions, except that coverslips were washed with Wash Buffer A for 3x instead of 2x and volumes used for probe incubation, ligation, and amplification were reduced by 25%. Coverslips were mounted on slides with Fluoroshield (Sigma-Aldrich F6182) with 1 μg/ml DAPI. Nuclei were imaged with an inverted spinning disk confocal (Andor Dragonfly/Leica) with a 100x oil objective. Only PLA foci in EdU positive cells were quantified using the Find Maxima feature in ImageJ.

### TCGA data analysis and visualization

R (version 4.1.0) and R package ComplexHeatmap (https://doi.org/10.1093/bioinformatics/btw313) was used for the data analysis and visualization for this project. Both the RNA-seq (FPKM-UQ), somatic variant calling, copy number variation, and clinical information data were downloaded through the GDC portal (https://portal.gdc.cancer.gov/). Breast PAM50 subtypes as well as important gene mutation, copy number alteration and genomic signatures were labeled at the top of the heatmap. Replication Stress (RS) high tumors are defined as whether a tumor carries at least one of the following mutations: CCNE1 amplification, MYC and MYCL1 amplification, KRAS amplification, NF1 mutation, RB1 two-copy loss, CDKN2A two-copy loss, ERBB2 amplification.^91^ Homologous recombination deficiency (HRD) is defined as the unweighted sum of three signatures: Number of telomeric Allelic Imbalances (NtAI), Large-scale State Transitions (LST), and loss of heterozygosity (LOH).^92^ CA20 score was defined as the sum of the expression of 20 genes (log2 median centered). These genes include: AURKA, CCNA2, CCND1, CCNE2, CDK1, CEP63, CEP152, E2F1, E2F2, LMO4, MDM2, MYCN, NDRG1, NEK2, PIN1, PLK1, PLK4, SASS6, STIL and TUBG1.^93^ To generate the heatmap, the gene expression matrix was first transformed using a log2 scale and then standardized for each gene. This standardized matrix was then used with the Heatmap function from the ComplexHeatmap package to create the heatmap, employing hierarchical clustering based on Euclidean distance. The Pearson’s correlation test was performed by the default function in R cor.test() on the log2-transformed gene FPKM expression value.

### Luciferase assay

*RNF25^+/+^* and *RNF25^-/-^* Pa02C cells were lipofectamine transfected with 100ng NF-κB luciferase reporter plasmid (a kind gift from Dr. Albert Baldwin at UNC Chapel Hill) and 1ng pRL (Renilla Luciferase Control Reporter, Promega E2241) for 24 h with media change after 6 h of transfection. Cell monolayers were further incubated in fresh media containing WEE1i for 24 h. Cells were harvested and luciferase activities were measured by using the dual luciferase reporter assay (Promega E1910) according to the manufacturer’s protocol.

For experiments with ectopic expression of HA-RNF25, *RNF25^+/+^*Pa02C cells were lipofectamine transfected with pRL/NFκB reporter plasmids for 6 h followed by RNF25 adenovirus infection at 1×10^11^ pfu/ml for 24 h. Cell monolayers were then incubated in media with WEE1i for about 18 h before harvesting for luminescence measurements.

### Statistical analysis

Graphs were generated and statistical analyses were performed using GraphPad Prism v9.4.1 and v10.1.1 (GraphPad Software). All statistical tests performed are described in Figure Legends for each respective experiment.

## Resource Availability

### Lead Contact

Requests for additional information and resources may be directed to Cyrus Vaziri. This work is a preprint and has not been peer reviewed.

### Materials Availability

Reagents generated in this study are available upon request.

### Data and Code Availability

- The mass spectrometry proteomics data have been deposited to the ProteomeXchange Consortium via the PRIDE partner repository with the dataset identifier PXD056698. This data will be publicly available as of the date of publication.
- CRISPR-Cas9 genetic screen data have been deposited at Zenodo and are publicly available as of the date of publication at (10.5281/zenodo.13930006).
- This paper does not report original code.

Any additional information required to reanalyze the data reported in this paper is available from the lead contact upon request.

## Acknowledgments

We thank Dr. Grant Stewart and Dr. Sati Jhujh for advice with DNA fiber assays and Dr. Pablo Ariel for his guidance in microscopy image acquisition. This work was supported by R01 ES029079 and CA215347 from the National Institutes of Health to CV. LFC and GND were supported in part by a grant from the National Institute of General Medical Sciences (5T32GM135128). This research is based in part upon work conducted using the UNC Proteomics Core Facility, which is supported in part by NCI Center Core Support Grant (2P30CA016086-45) to the UNC Lineberger Comprehensive Cancer Center. The Andor Dragonfly microscope was funded with support from National Institutes of Health grant (S10OD030223). The UNC Flow Cytometry Core Facility (RRID:SCR_019170) is supported in part by P30 CA016086 Cancer Center Core Support Grant to the UNC Lineberger Comprehensive Cancer Center. The Thermo Fisher Attune NxT was supported in part by the North Carolina Biotech Center Institutional Support Grant 2017-IDG-1025 and by the National Institutes of Health 1UM2AI30836-01. The Becton Dickinson FACSAria II was supported in part by the North Carolina Biotech Center Institutional Support Grant 2012-IDG-1006. The content is solely the responsibility of the authors and does not necessarily represent the official views of the National Institutes of Health.

## Author Contributions

All authors participated in designing or discussing experiments. LFC, DJ, GND, XZ, YY, CAM, NKB, and TSW performed experiments. LFC, DJ, and GND prepared figures. LFC and CV wrote the paper, with support from DJ and GND. DW, LEH, JB, and CV supervised experiments and analysis. CV conceived the project and provided funding. All authors reviewed and edited the paper.

