## Supplementary Figures 1-4, Supplementary Tables 1-4 for "The RING Finger E3 Ligase RNF25 Protects DNA Replication Forks Independently of its Canonical Roles in Ubiquitin Signaling"

### Supplementary Fig. 1

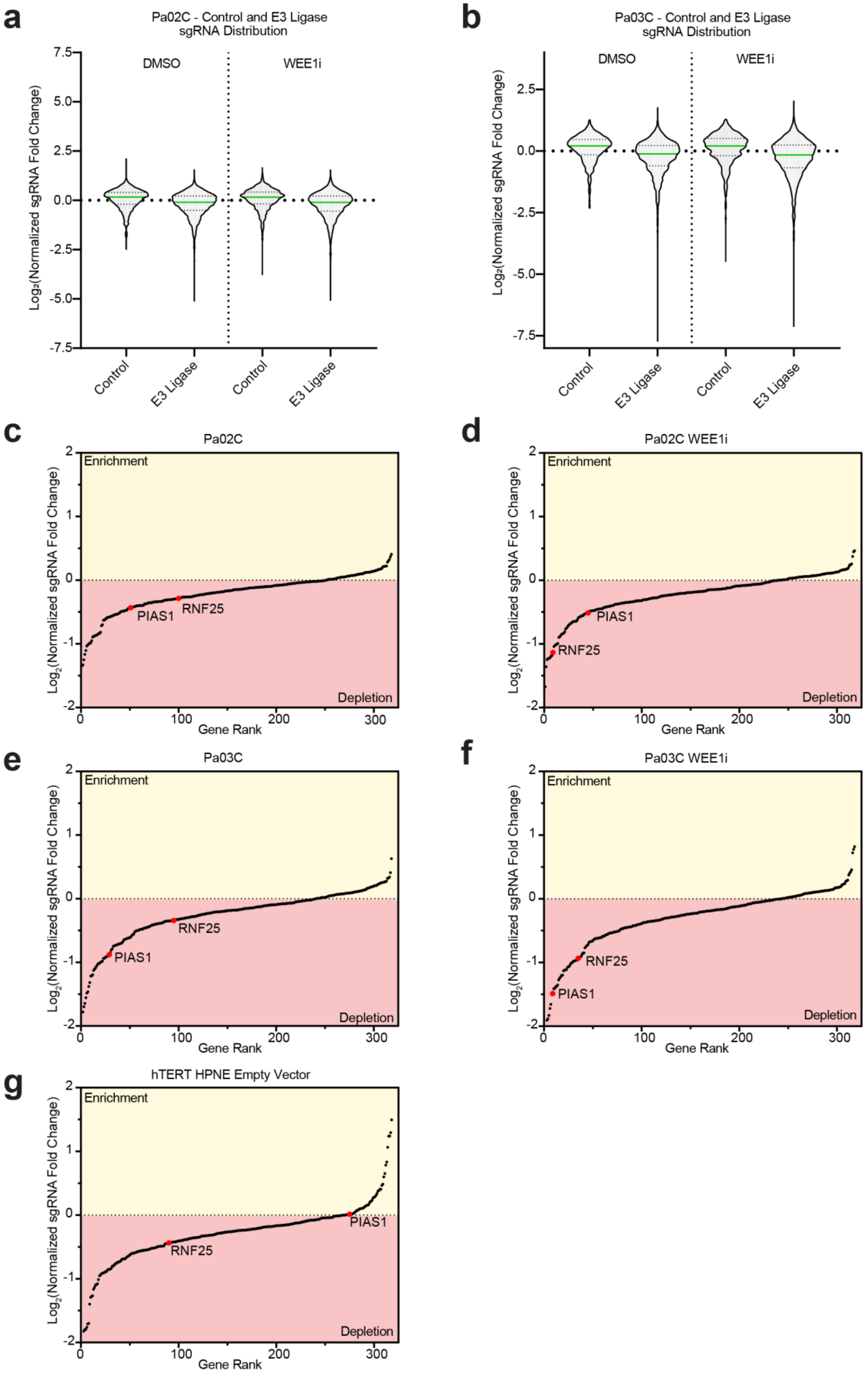

**Supplementary Fig. 1: RNF25 and PIAS1 show increased gene ranks upon WEE1 inhibitor treatment in PDAC cell lines.**

**a, b** Relative distribution of control non-targeting sgRNAs and RING-finger E3 ligase-directed sgRNAs in Pa02C and Pa03C cells after 20 population doublings ± WEE1 inhibitor treatment.

**c, d, e, f** Gene rank plots showing relative enrichment or depletion of RING-finger E3 ligase-directed sgRNAs in Pa02C and Pa03C cells after 20 population doublings ± WEE1 inhibitor treatment.

**g** Gene rank plot showing relative enrichment or depletion of RING-finger E3 ligase-directed sgRNAs after 20 population doublings in hTERT-immortalized normal human pancreatic epithelial cells (HPNE).

All data including sgRNA distribution violin and gene rank plots were gathered from at least three biological replicates.

### Supplementary Fig. 2

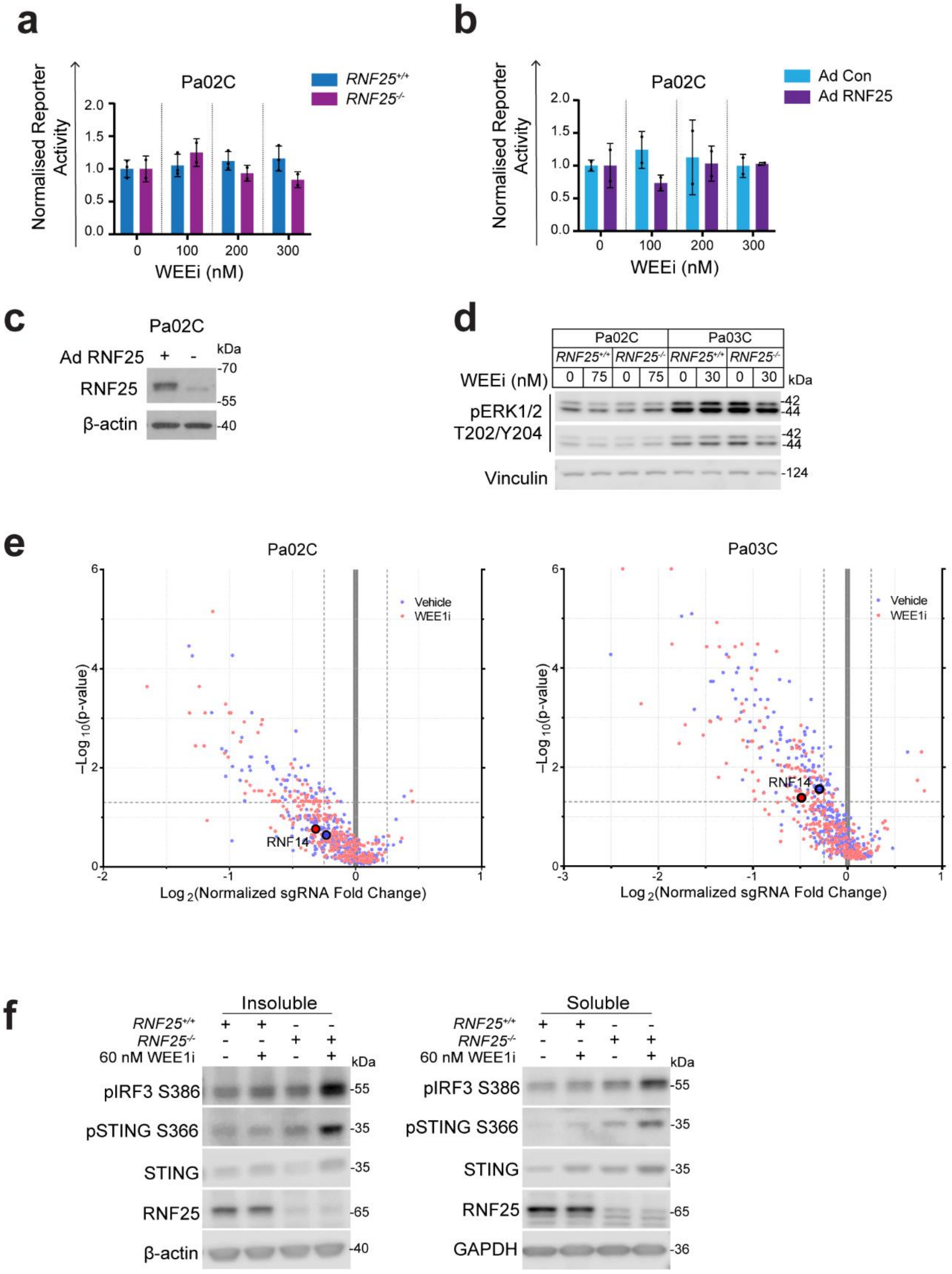

**Supplementary Fig. 2: RNF25-dependent WEE1i-tolerance is unrelated to NF-κB, ERK, or translation quality control.**

**a** Effect of WEE1i on expression of NFκB-responsive luciferase reporter activity in *RNF25*$^{+/+}$ and *RNF25*$^{-/-}$ Pa02C cells.

**b** Effect of adenovirally-overexpressed HA-RNF25 on expression of NFκB-responsive luciferase reporter activity in *RNF25*$^{+/+}$ Pa02C cells.

**c** Immunoblots showing levels of RNF25 in *RNF25*$^{+/+}$ and HA-RNF25-overexpressing *RNF25*$^{+/+}$ Pa02C cells.

**d** Immunoblot showing effect of WEE1i treatment on levels of active T202/Y204 dually-phosphorylated pERK in *RNF25*$^{+/+}$ and *RNF25*$^{-/-}$ Pa02C and Pa03C cells after conditional treatment with WEE1i.

**e** Results of CRISPR screens showing dropout of RNF14-directed sgRNAs in relation to all other library sgRNAs in vehicle (DMSO)- and WEE1i-treated Pa02C and Pa03C cells. $Log_2$-transformed sgRNA abundance scores and statistical significance were calculated using Völundr.

**f** Immunoblot showing effect of WEE1i treatment on the indicated protein markers of the cGAS-STING pathway in *RNF25*$^{+/+}$ and *RNF25*$^{-/-}$ Pa03C cells.

All reporter activity data (a, b) represent mean ± SD from duplicate dishes and are representative of two biological replicates. Statistics: two-way ANOVA with Sidak's multiple comparisons test.

Supplementary Fig. 3

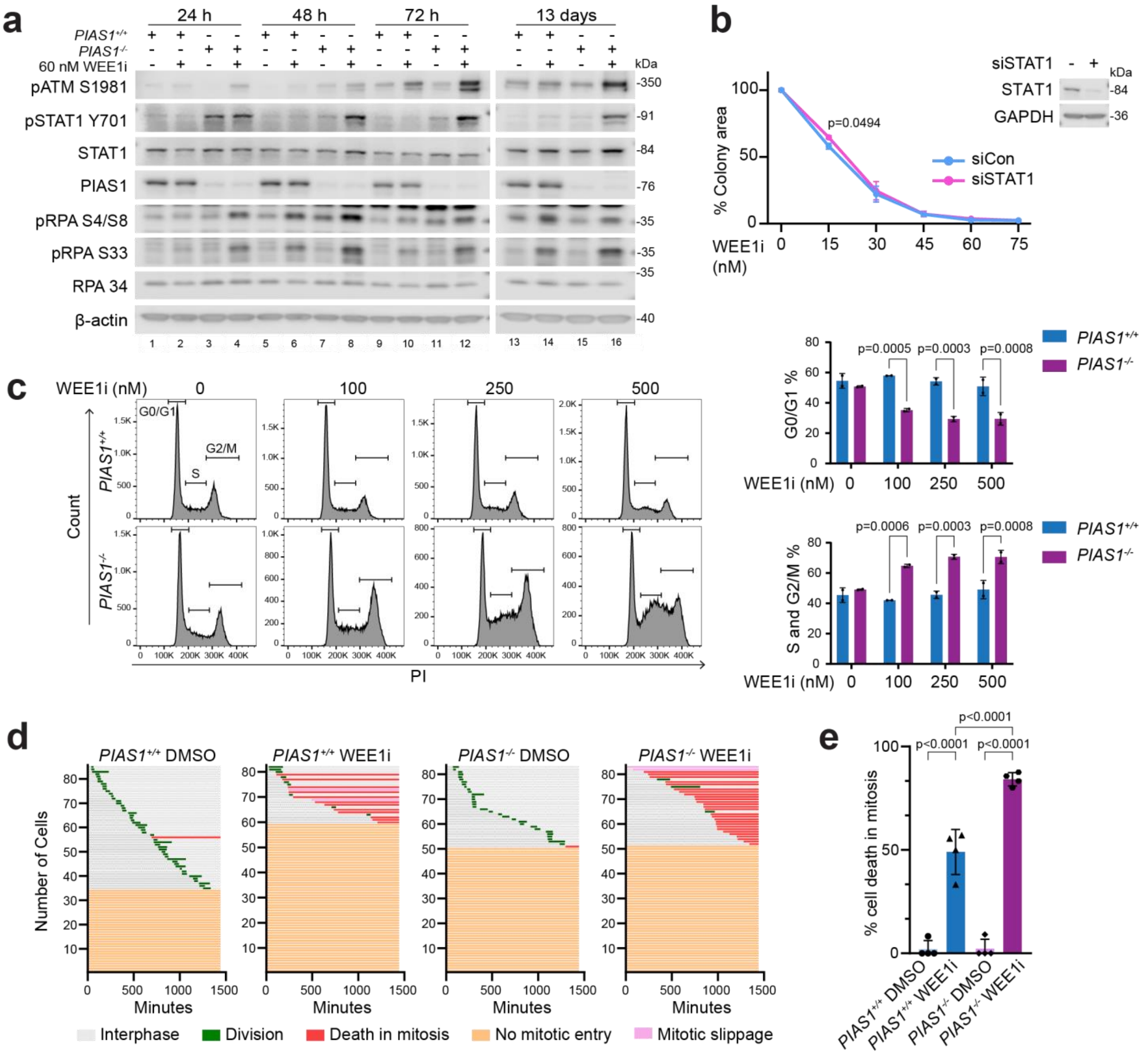


**Supplementary Fig. 3: PIAS1 promotes S-phase and G2/M progression in WEE1i-treated PDAC cells.**

**a** Immunoblot showing effect of WEE1i treatment on levels of the indicated DNA damage markers in *PIAS1*$^{+/+}$ and *PIAS1*$^{-/-}$ Pa03C cells at the indicated timepoints.

**b** Clonogenic survival assay showing effect of STAT1 siRNA on WEE1i-sensitivity of *PIAS1*$^{-/-}$ Pa03C cells. Statistics were analyzed using a two-way ANOVA with Sidak's multiple comparisons test.

**c** FACS analyses showing effect of WEE1i treatment on cell cycle profiles of *PIAS1*$^{+/+}$ and *PIAS1*$^{-/-}$ Pa03C cells. The cell cycle analyses for the *PIAS1*$^{+/+}$ cells shown here are the same profiles shown for *RNF25*$^{+/+}$ cells in Fig. 2b. These Pa03C cells were transduced with non-targeting sgRNAs and are wild-type for both RNF25 and PIAS1. All the cell cycle analyses shown in Fig. 2b and Supplementary Fig. 3c were performed at the same time. These data represent mean ± SD from duplicate dishes. All data are from a representative experiment that yielded similar results on two occasions. Statistics: two-way ANOVA with Sidak's multiple comparisons test.

**d** Cell fate maps showing effect of WEE1i-treatment on interphase and mitotic progression of *PIAS1*$^{+/+}$ and *PIAS1*$^{-/-}$ Pa03C cells, as determined using live cell imaging experiments.

**e** Quantification of results from (d) showing effect of WEE1i-treatment on mitotic cell death in *PIAS1*$^{+/+}$ and *PIAS1*$^{-/-}$ Pa03C cells. All data represent mean ± SD from four focus locations. Ordinary one-way ANOVA with Tukey's multiple comparisons test was performed to analyze statistical significance.

### Supplementary Fig. 4

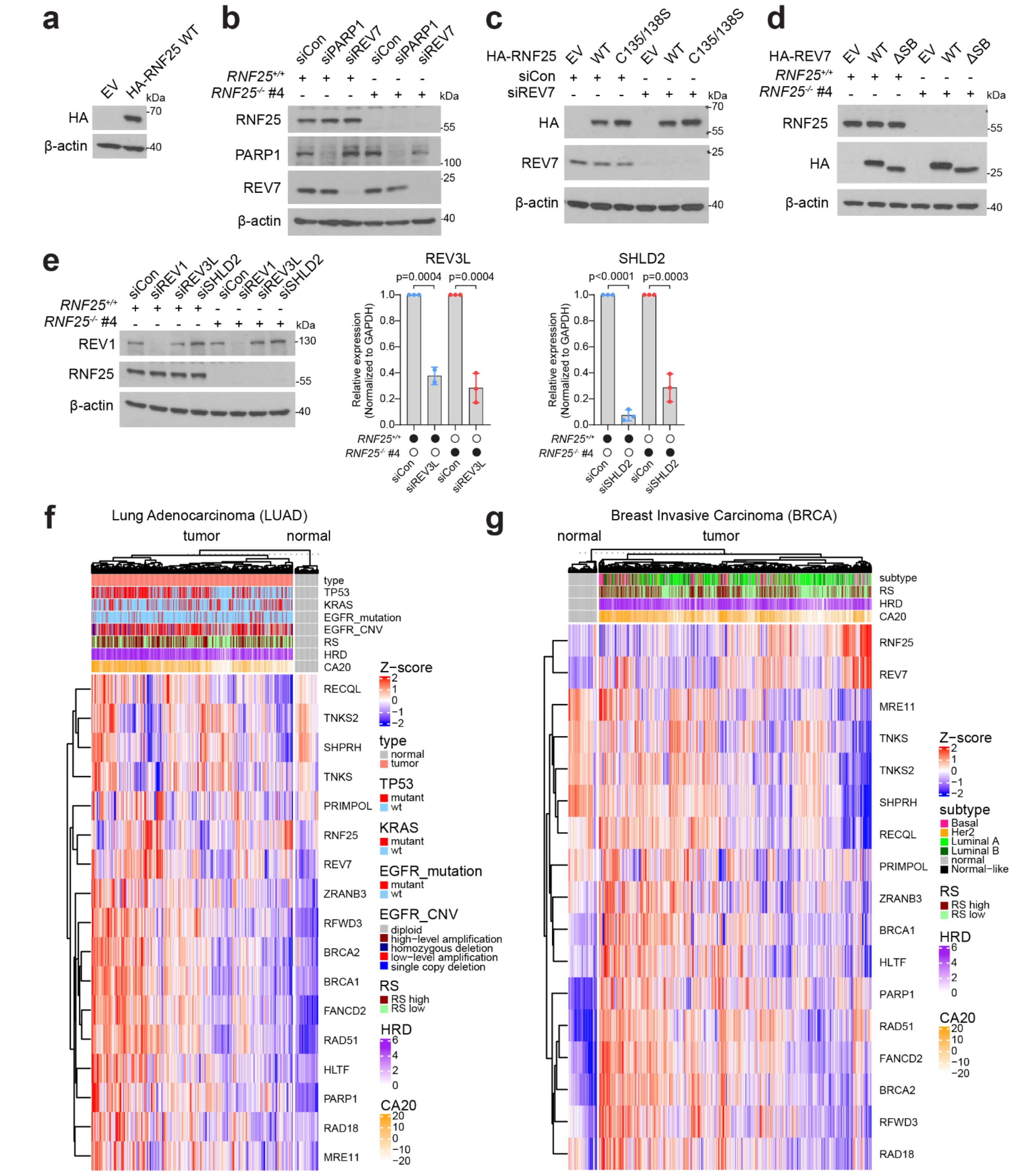


**h**

| TCGA dataset | Pearson's Correlation Coefficient (r) | p-value |
|---|---|---|
| PDAC | 0.51 | <0.001 |
| LUAD | 0.50 | <0.001 |
| BRCA | 0.36 | <0.001 |

**Supplementary Fig. 4: RNF25 and REV7 expression are significantly correlated.**

**a** Immunoblots showing validation of ectopic HA-RNF25 expression in the SIRF assay shown in Fig. 4d.

**b** Immunoblots showing validation of REV7 and PARP1 knockdown in the DNA fiber assay shown in Fig. 6a, b.

**c** Immunoblots validating ectopic expression of HA-RNF25 WT and HA-RNF25 C135/138S, and knockdown of REV7 in the SIRF assay shown in Fig. 6d.

**d** Immunoblots validating ectopic expression HA-REV7 WT and HA-REV7ΔSB mutant in the SIRF assay shown in Fig. 6e.

**e** Immunoblot validating knockdown of REV1 protein (left panel) and results of qPCR analysis validating knockdown of REV3L and SHLD2 mRNAs in the DNA fiber assay shown in Fig. 7a. Statistics for qPCR analysis were calculated using a two-tailed Student's t-test.

**f, g** Heatmaps depicting relative mRNA expression of DNA replication and DNA damage repair factors in LUAD (f) and BRCA (g) patient tumor samples from TCGA. Supervised clustering was performed to separate normal and tumor samples. The heatmap in panel (f) shows mutation status of genes that are commonly altered in LUAD. Other classifiers shown include tumor subtype (shown for BRCA), and genome maintenance related features (RS – Replication Stress signature, HRD – Homologous Recombination Deficiency score, CA20 – Centrosome Amplification signature).

**h** Results of Pearson's Correlation Coefficient test for RNF25 and REV7 expression in the indicated TCGA datasets. The coefficient r value and p-value are indicated.

#### Supplementary Table 1

| Gene | Log2(normalized sgRNA fold change) | Corrected p-value | –Log10 p-value (K-S test) |
|---|---|---|---|
| UBE2I | -1.34 | 5.30E-05 | 4.276 |
| TRAIP | -1.33 | 3.50E-04 | 3.456 |
| RNF113A | -1.25 | 2.32E-03 | 2.634 |
| TRIM28 | -1.17 | 6.54E-04 | 3.184 |
| RBBP6 | -1.10 | 3.58E-03 | 2.447 |
| MSL2 | -1.03 | 3.00E-03 | 2.522 |
| NSMCE1 | -1.02 | 8.88E-03 | 2.051 |
| RANBP2 | -1.01 | 1.70E-02 | 1.769 |
| COP1 | -0.99 | 2.20E-05 | 4.658 |
| TRIM61 | -0.99 | 3.19E-01 | 0.496 |
| RNF4 | -0.95 | 5.76E-03 | 2.240 |
| TRIM64B | -0.89 | 1.67E-03 | 2.779 |
| RNF168 | -0.88 | 2.46E-02 | 1.608 |
| BARD1 | -0.88 | 5.76E-03 | 2.240 |
| PCGF7P | -0.87 | 4.43E-02 | 1.353 |
| NSMCE2 | -0.86 | 6.54E-04 | 3.184 |
| RCHY1 | -0.86 | 1.40E-02 | 1.853 |
| MARCH6 | -0.84 | 5.76E-03 | 2.240 |
| TRAF2 | -0.84 | 5.76E-03 | 2.240 |
| RNF8 | -0.82 | 5.76E-03 | 2.240 |

#### Supplementary Table 2

| Gene | Log2(normalized sgRNA fold change) | Corrected p-value | –Log10 p-value (K-S test) |
|---|---|---|---|
| UBE2I | -1.67 | 2.77E-04 | 3.558 |
| NSMCE1 | -1.36 | 2.77E-04 | 3.558 |
| MSL2 | -1.25 | 2.77E-04 | 3.558 |
| TRAIP | -1.23 | 5.97E-04 | 3.224 |
| TRIM28 | -1.23 | 9.41E-04 | 3.026 |
| RNF8 | -1.22 | 4.38E-03 | 2.359 |
| RNF113A | -1.19 | 9.41E-04 | 3.026 |
| TRIM61 | -1.18 | 1.66E-01 | 0.780 |
| RNF25 | -1.13 | 3.00E-06 | 5.523 |
| NSMCE2 | -1.04 | 7.75E-04 | 3.111 |
| RNF4 | -1.02 | 5.97E-04 | 3.224 |
| TRIM49 | -1.01 | 1.59E-02 | 1.798 |
| COP1 | -1.00 | 3.33E-04 | 3.478 |
| RNF168 | -0.99 | 7.75E-04 | 3.111 |
| MARCH6 | -0.90 | 9.41E-04 | 3.026 |
| RANBP2 | -0.89 | 5.70E-03 | 2.244 |
| TRIM64B | -0.87 | 3.35E-02 | 1.475 |
| RBBP6 | -0.85 | 2.04E-02 | 1.691 |
| ANAPC11 | -0.84 | 2.40E-02 | 1.619 |
| TRAF2 | -0.83 | 1.69E-02 | 1.772 |

**Supplementary Table 3**

| Gene | Log2(normalized sgRNA fold change) | Corrected p-value | –Log10 p-value (K-S test) |
|---|---|---|---|
| UBE2I | -2.53 | 6.70E-05 | 4.174 |
| TRIM28 | -1.78 | 1.00E-05 | 5.000 |
| RNF4 | -1.70 | 3.00E-06 | 5.523 |
| MIB1 | -1.64 | 5.19E-04 | 3.285 |
| RNF113A | -1.56 | 8.60E-05 | 4.066 |
| NSMCE1 | -1.48 | 1.81E-04 | 3.742 |
| NSMCE2 | -1.46 | 2.59E-04 | 3.587 |
| TRIM61 | -1.38 | 1.60E-03 | 2.796 |
| BARD1 | -1.32 | 3.60E-05 | 4.444 |
| RNF216 | -1.21 | 2.60E-04 | 3.585 |
| BRCA1 | -1.21 | 5.19E-04 | 3.285 |
| TRAIP | -1.19 | 1.95E-04 | 3.710 |
| TRAF7 | -1.13 | 7.50E-05 | 4.125 |
| RNF8 | -1.10 | 7.93E-04 | 3.101 |
| ANAPC11 | -1.08 | 8.45E-03 | 2.073 |
| TRIM11 | -1.06 | 3.61E-03 | 2.443 |
| SENP6 | -1.03 | 2.31E-03 | 2.637 |
| ZNRF2 | -1.02 | 7.06E-04 | 3.151 |
| MNAT1 | -1.02 | 3.64E-04 | 3.439 |
| MARCH6 | -1.00 | 3.64E-04 | 3.439 |

**Supplementary Table 4**

| Gene | Log2(normalized sgRNA fold change) | Corrected p-value | –Log10 p-value (K-S test) |
|---|---|---|---|
| UBE2I | -2.40 | 0.00E+00 | 6.000 |
| MIB1 | -2.21 | 3.14E-04 | 3.503 |
| NSMCE1 | -1.91 | 2.00E-06 | 5.699 |
| TRIM28 | -1.88 | 4.40E-05 | 4.357 |
| NSMCE2 | -1.83 | 3.66E-03 | 2.437 |
| TRIM11 | -1.72 | 2.00E-03 | 2.700 |
| RNF113A | -1.66 | 3.97E-04 | 3.401 |
| ANAPC11 | -1.50 | 5.57E-03 | 2.254 |
| PIAS1 | -1.49 | 4.40E-05 | 4.357 |
| TRAF7 | -1.41 | 1.49E-03 | 2.827 |
| RNF4 | -1.41 | 4.30E-05 | 4.367 |
| RNF216 | -1.38 | 1.90E-05 | 4.721 |
| TRIM61 | -1.37 | 9.25E-03 | 2.034 |
| RNF8 | -1.35 | 1.29E-03 | 2.890 |
| MARCH6 | -1.27 | 5.90E-05 | 4.229 |
| SENP5 | -1.25 | 1.29E-02 | 1.890 |
| TRAIP | -1.23 | 3.14E-04 | 3.503 |
| RNF126 | -1.20 | 1.29E-03 | 2.890 |
| COP1 | -1.20 | 6.81E-04 | 3.167 |
| BARD1 | -1.13 | 3.14E-04 | 3.503 |

**Supplementary Table 1-4: Top 20 depleted RING-finger E3 ubiquitin/SUMO ligase sgRNAs in Pa02C and Pa03C CRISPR Cas9 screens (Related to Fig. 1).**

CRISPR screen results showing relative sgRNA depletion or enrichment in Pa02C/Pa03C control (Supplementary Table 1, 3) and WEE1i-treated groups (Supplementary Table 2, 4). Kolmogorov–Smirnov test statistic and its corrected p-value are shown compared to the distribution of 1,000 control sgRNAs.
